# Membrane binding of a cyanobacterial ESCRT-III protein crucially involves the helix α1-3 hairpin conserved in all superfamily members

**DOI:** 10.1101/2025.09.05.674404

**Authors:** Lukas Schlösser, Mirka Kutzner, Nadja Hellmann, Denis Kiesewetter, Julia Bieber, Ndjali Quarta, Xingwu Ge, Tom Goetze, Benedikt Junglas, Fumiki Matsumura, Mischa Bonn, Frauke Gräter, Carsten Sachse, Lu-Ning Liu, Carla Schmidt, Camilo Aponte-Santamaría, Dirk Schneider

## Abstract

IM30, the inner membrane-associated protein of 30 kDa (also known as Vipp1) is essential for thylakoid membrane biogenesis and/or maintenance in chloroplasts and cyanobacteria. IM30 and its bacterial homolog PspA belong to the ESCRT-III superfamily, proteins previously thought to be restricted to eukaryotes and archaea. Despite low sequence similarity, IM30 shares key structural and functional features with eukaryotic ESCRT-IIIs, including a conserved α1–α2 helical hairpin core and the ability to form oligomeric barrel- or rod assemblies that mediate membrane remodeling. Using IM30 variants, we now show that initial membrane recruitment of IM30 is driven by electrostatic interactions between the positively charged α1–α3 helical hairpin and negatively charged lipid surfaces, paralleling the role of charged helical regions in some eukaryotic ESCRT-IIIs. This likely initiates lateral assembly of IM30 into higher-order barrel or rod structures on the membrane. Once assembled, α0 helices within these oligomers engage and stabilize internalized membrane tubules, mirroring membrane interaction strategies of eukaryotic ESCRT-IIIs, which use both N-terminal sequences and charged residues on α1/α2. Thus, our findings demonstrate a conserved membrane binding and remodeling mechanism across the ESCRT-III superfamily, underscoring an evolutionary link in membrane dynamics between pro- and eukaryotes.

**Significance:** IM30, a membrane-associated protein found in cyanobacteria and chloroplasts, along with its bacterial homolog PspA, belongs to the ESCRT-III superfamily. Despite low sequence conservation, these proteins share structural and functional features with eukaryotic ESCRT-III proteins. We show that IM30 binds membranes via a conserved structural motif, followed by lateral assembly into higher-order complexes. This supports a mechanism of membrane remodeling that is conserved in prokaryotic and eukaryotic members of the ESCRT-III superfamily.

## Introduction

Photosynthesis, the process by which plants, algae, and cyanobacteria convert light energy into chemical energy, involves the initial capture of light to drive photosynthetic electron transport within thylakoid membranes (TMs), a specialized intracellular membrane system (Johnson, 2025; Perez-Boerema et al., 2024; Pribil et al., 2014). In plants and algae, TMs are housed within chloroplasts, while cyanobacteria possess an analogous extra internal TM system, reflecting their evolutionary ties to modern chloroplasts (Mullineaux and Liu, 2020; Ostermeier et al., 2024; Perez-Boerema et al., 2024). Despite their importance, the biogenesis, dynamics, and maintenance of TM systems remain poorly understood in both chloroplasts and cyanobacteria (Ostermeier et al., 2024).

A protein crucially involved in the biogenesis and maintenance of TMs in chloroplasts and cyanobacteria is the *inner membrane-associated protein of 30 kDa* (IM30), also known as the *vesicle-inducing protein in plastids 1* (Vipp1) (Kroll et al., 2001; Vothknecht et al., 2012; Westphal et al., 2001). In chloroplasts, IM30 exhibits a dual localization, being distributed within the stroma and associated with both the inner envelope membrane and TMs (Kroll et al., 2001; Li et al., 1994). Similarly, in cyanobacteria, IM30 exhibits dynamic localization, being found throughout the cytoplasm and bound to both the cytoplasmic membrane and TMs (Bryan et al., 2014; Fuhrmann et al., 2009a; Gutu et al., 2018).

Studies of *vipp1* knock-down and knock-out mutants of *Arabidopsis thaliana* have shown that the protein is essential for the biogenesis and/or maintenance of chloroplast TMs and inner envelope membranes (Aseeva et al., 2007; Kroll et al., 2001; Zhang et al., 2012; Zhang et al., 2016a). The deletion of *im30* is lethal in cyanobacteria, underscoring its indispensable role in these organisms. Cyanobacterial cells with reduced IM30 levels display impaired TM morphologies and compromised functional integrity (Fuhrmann et al., 2009b; Gao and Xu, 2009; Westphal et al., 2001).

A structural relationship between IM30 and the bacterial *phage shock protein A* (PspA) has been recognized early on, suggesting that the *im30* gene evolved from a *pspA* gene duplication event (Bultema et al., 2010; Kroll et al., 2001; Vothknecht et al., 2012; Westphal et al., 2001). Notably, *im30* expression can functionally complement a *pspA* deletion in *Escherichia coli* (DeLisa et al., 2004; Zhang et al., 2012). While IM30 is present in both cyanobacteria and chloroplasts, PspA is restricted to certain bacteria, including cyanobacteria, and is absent in chloroplasts (Popp et al., 2022; Ravi et al., 2024; Vothknecht et al., 2012). In PspA-containing bacteria, the protein’s primary function appears to be the maintenance and repair of the cytoplasmic membrane, although it is not essential for viability (Darwin, 2005; Manganelli and Gennaro, 2017).

Recent structural analyses have categorized IM30 and PspA as members of the *endosomal sorting complex required for transport III* (ESCRT-III) superfamily, a class of proteins previously thought to be exclusive to eukaryotes and archaea (Gupta et al., 2021; Junglas et al., 2021; Liu et al., 2021). Despite low sequence identity with their eukaryotic counterparts, prokaryotic and eukaryotic ESCRT-III members share striking similarities in secondary and tertiary structure, as well as membrane-remodeling activity (McCullough and Sundquist, 2025; Nachmias et al., 2025; Pfitzner et al., 2021; Schlosser et al., 2023). A defining feature of ESCRT-III proteins is the presence of at least five α-helices, with the long helices α1 and α2 forming a hairpin structure, the structural core of all ESCRT-III superfamily members (McCullough and Sundquist, 2020; Schlosser et al., 2023). Additionally, all ESCRT-III superfamily members exhibit an intrinsic propensity to form large oligomeric assemblies, with bacterial proteins forming homooligomeric structures and eukaryotic counterparts typically assembling into heterooligomeric complexes (McCullough et al., 2018; Schlosser et al., 2023).

An IM30 monomer comprises seven α-helices (α0–α6), with the first six helices (α0–α5) forming a structural domain homologous to PspA, as revealed by recent cryo-electron microscopy (cryo-EM) studies (Gupta et al., 2021; Junglas et al., 2025; Liu et al., 2021). The IM30-specific helix α6 appears to play a crucial role in IM30’s *in vivo* activity, although its exact physiological function remains unclear (Hennig et al., 2017; Ma et al., 2025; Zhang et al., 2016b). The core of IM30 is defined by a helical hairpin formed by α1–3, with α3 being a direct extension of α2 (Gupta et al., 2021; Liu et al., 2021). In oligomeric assemblies, helices α1–6 adopt α-helical conformations, but upon disassembly, only the α1–3 hairpin retains its α-helical structure, while α0 and α4–6 unfold (Junglas et al., 2020; Quarta et al., 2024). An engineered IM30 variant (IM30*) with disrupted intersubunit contacts, yet preserved helical propensity, is incapable of forming large oligomers, thereby mimicking the disassembled IM30 state (Heidrich et al., 2016; Junglas et al., 2020; Quarta et al., 2024).

In the absence of membranes, wt IM30 self-assembles into diverse homo-oligomeric barrel structures, with currently described internal symmetries ranging from 7–22 in *Synechocystis* (Fuhrmann et al., 2009a; Gupta et al., 2021; Junglas et al., 2025; Saur et al., 2017) and 11–17 in *Nostoc punctiforme* (Liu et al., 2021). Each monomer interacts with multiple neighboring subunits in both axial directions, forming barrel-like structures and higher-order rod-shaped complexes with masses of several MDa (Gupta et al., 2021; Junglas et al., 2025; Liu et al., 2021; Schlosser et al., 2023). The individual helices of IM30 monomers are connected by six hinge regions (hinge 0: α0 – α1; hinge 1: α2 – α 3; hinge 2: α3 – α 4; hinge 3: α4 – α 5; hinge 4: α5 – α6), enabling flexibility and variable ring and rod diameters, as well as dome-like architectures of the barrel assemblies.

IM30 oligomers bind negatively charged membrane surfaces (Heidrich et al., 2016; Hennig et al., 2015; Theis et al., 2019; Thurotte and Schneider, 2019), forming carpets and spirals upon membrane binding, which likely involves disassembly of the homooligomeric barrel and rod structures observed in solution (Junglas et al., 2022; Junglas et al., 2025; Junglas et al., 2020; Naskar et al., 2025; Pan et al., 2024). Monomeric IM30 can oligomerize to form large IM30 assemblies that internalize tubulated membranes, mediated by the N-terminal amphiphilic helix α0, which partially embeds into the lipid bilayer (Junglas et al., 2025). Similar membrane internalization into PspA rods has been observed, with α0 removal abolishing this activity (Hudina et al., 2025). IM30 and PspA monomers likely assemble on membrane surfaces, resulting in the formation of barrel or rod structures, concurrently "sucking in" membranes to stabilize barrel and tubular membrane architectures (Gupta et al., 2021; Junglas et al., 2025; Junglas et al., 2020; Liu et al., 2021; Siebenaller et al., 2019). Recently, it has been suggested that IM30 and PspA barrels and/or rods bind membranes in their inner barrel/rod lumen via helix α0 (Gupta et al., 2021; Hudina et al., 2025; Junglas et al., 2025; Liu et al., 2021). Yet, the IM30* variant, lacking the intrinsic oligomerization propensity, still binds well to membranes, suggesting that protein oligomerization is no prerequisite for membrane binding *per se*, and multiple regions may contribute to membrane adhesion (Heidrich et al., 2016; Junglas et al., 2020; Nguyen et al., 2020). This latter notion is consistent with observations in eukaryotic ESCRT-III proteins, where membrane binding can be mediated by small amphipathic helical regions at the N-terminus (Buchkovich et al., 2013; McCullough et al., 2015) or positively charged residues on specific helices, as *e.g.* seen in human CHIMP-1B or yeast Snf7, which interacts with membranes via hydrophobic and positively charged residues located on helix α1 and α2 (Nguyen et al., 2020; Tang et al., 2015). Furthermore, the IM30 helix α6 has been implicated in membrane binding of *Synechocystis* IM30 (Hennig et al., 2017).

Using IM30 variants, we now demonstrate that the α1–α3 helical hairpin directly interacts with negatively charged membrane surfaces, while helix α0 enhances membrane binding. Based on our observations we propose that initial membrane recruitment of IM30 monomers is mediated primarily by the interactions of the α1–α3 hairpin with membranes, followed by lateral assembly on the membrane surface, akin to eukaryotic ESCRT-III proteins. This oligomerization results in the formation of spiral, barrel-, or rod-like structures. Within these assemblies, helix α0 then becomes the primary mediator of interactions with membranes mediating membrane tubulation and internalization. Together, our findings support a conserved membrane-binding and remodeling mechanism across the ESCRT-III superfamily, shared between prokaryotic IM30/PspA and their eukaryotic counterparts.

## Results

### IM30 binds to negatively charged membrane surfaces

Recently, membrane-binding of IM30 and its interaction with negatively charged membrane surfaces have been demonstrated via monitoring changes in lipid bilayer properties at increasing protein concentrations (Heidrich et al., 2016; Hennig et al., 2015). To assess membrane binding more directly, we now employed protein fluorescence emission spectroscopy, focusing on changes in IM30’s Trp fluorescence emission characteristics upon addition of purified proteins to negatively charged liposomes (Figure 1D). First, we determined the minimal fraction of negatively charged lipids required in net-neutral PC membranes to reliably detect membrane binding of IM30 wt via fluorescence spectroscopy (Figure 2 A, C).

**Figure 1:**
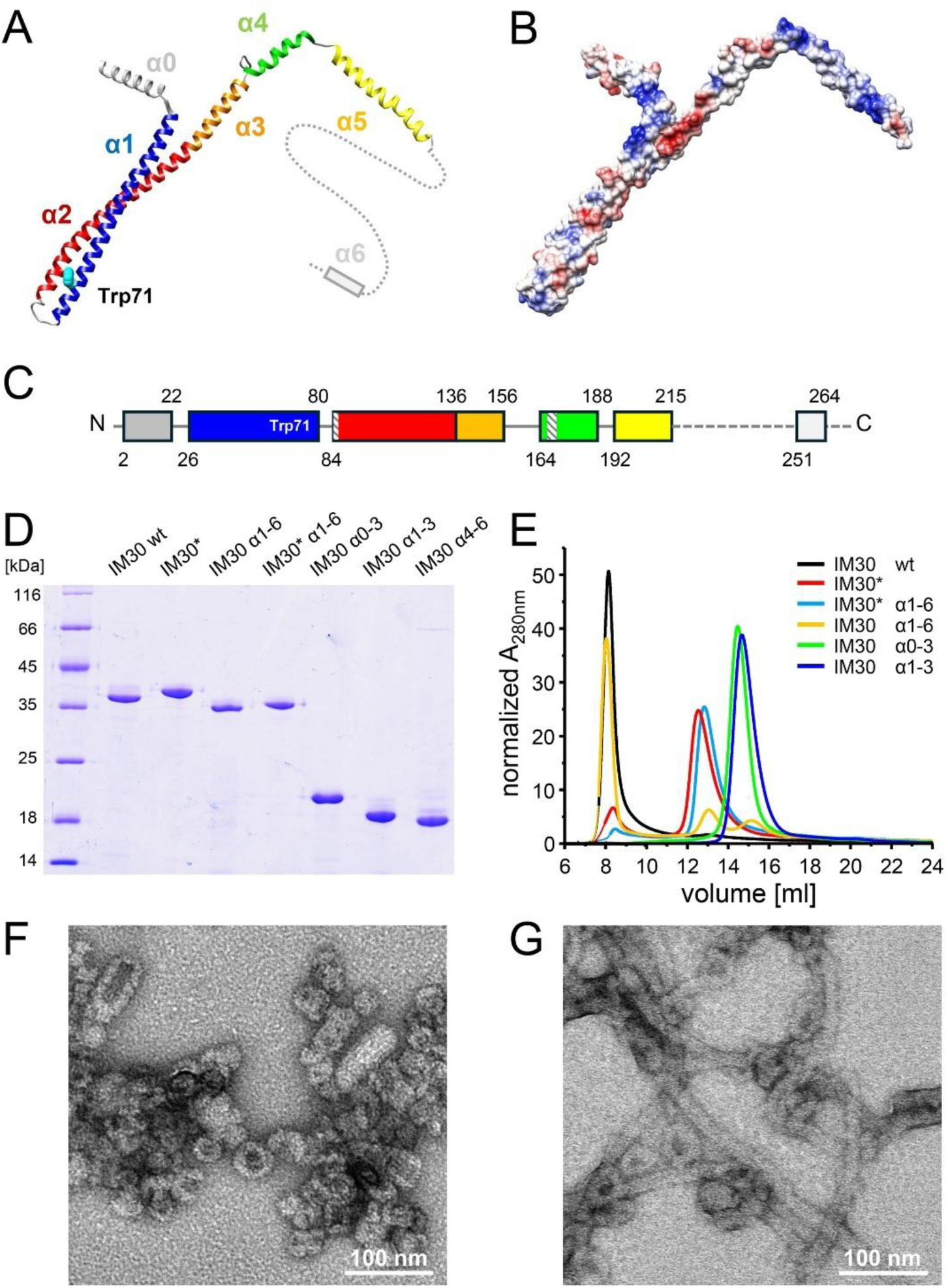
IM30 variants analyzed in the present study. (A, B) Structure (pdb: 703y) and (C) schematic illustration of IM30wt with helix numbering. The numbers give the beginnings and ends of the respective helices, as used for the expression of truncated IM30 variants. The dashed bars in (C) mark the regions mutated in IM30*. (B) Electrostatic surface potential of an α0-5 monomer (pdb: 703Y) with positively charged regions in red and negatively charged regions in blue. (D) SDS-PAGE analysis of the purified IM30 variants studied. (E) SEC profile of purified proteins normalized to the total area. Solely IM30 wt and IM30 α1-6 form large oligomeric structures. (F, G), EM of purified IM30 wt (F) and IM30 α1-6 (G) showing the formation of prototypical barrel structures, as well as stacked barrels, in the case of IM30 wt, whereas IM30 α1-6 forms elongated rod structures. Note that helix α6 is only predicted and has not been solved in any IM30 structure thus far.

**Figure 2:**
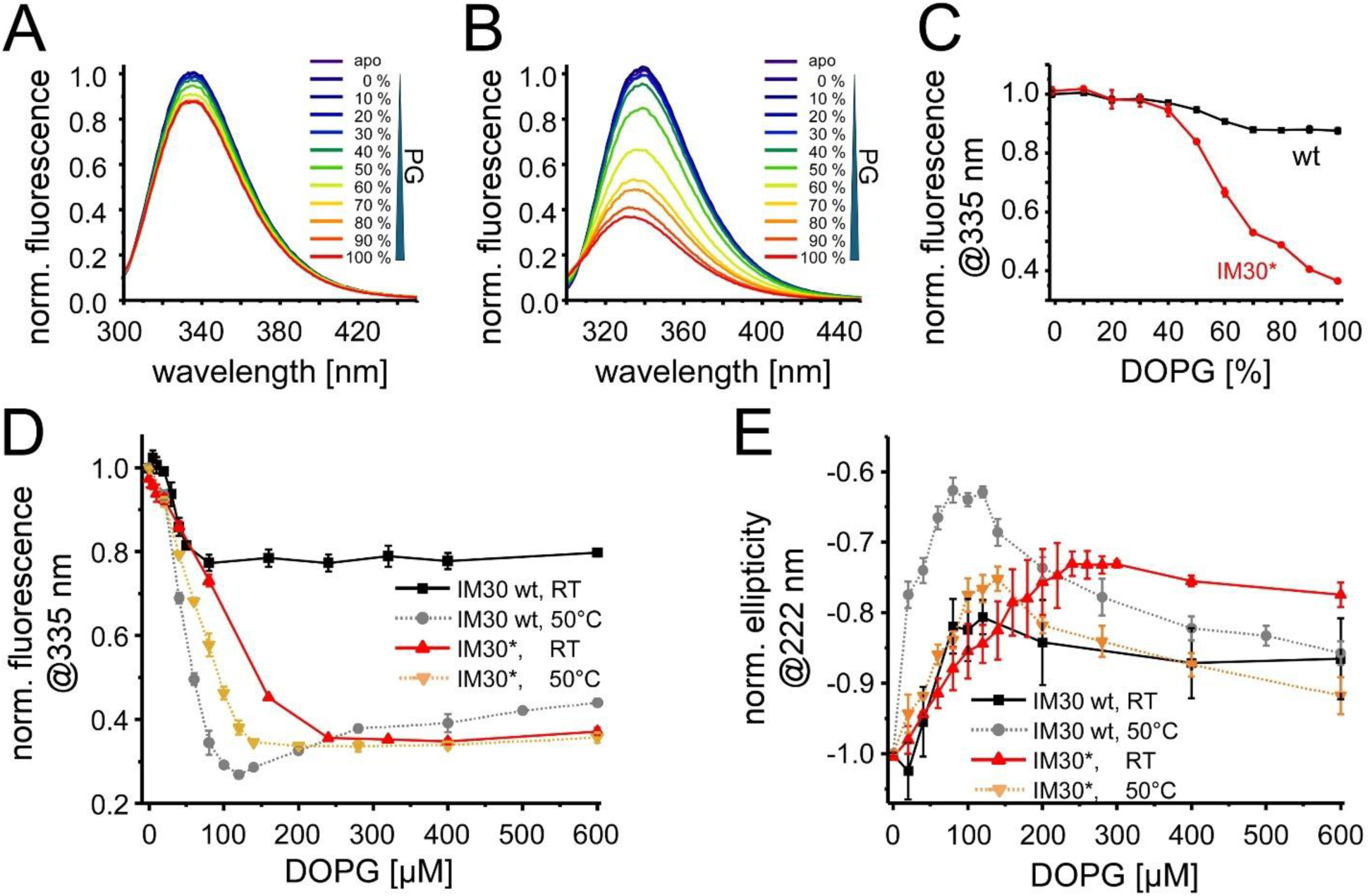
IM30(*) binding to PG-containing liposomes. (A, B) Normalized intrinsic protein fluorescence spectra of (A) IM30 wt and (B) IM30* in solution and after incubation for 2 hours with liposomes with increasing PG content. The fluorescence intensity decreases with increasing PG content (in a PC background). (C) Normalized fluorescence intensities of IM30 wt and IM30* at 335 nm (n=3, error bar=SD) at increasing PG contents. (D) Normalized fluorescence intensities of IM30 wt and IM30* at 335 nm determined at 25°C and 50°C in dependence of PG concentration. The protein fluorescence intensity decreased with increasing PG concentrations. The reduced fluorescence indicates membrane binding possibly also involving a rearrangement of the protein structure (n=3, error bars = SD). The corresponding spectra are shown in Figure S1. (E) Normalized ellipticity of IM30 wt and IM30* at 25 °C and 50 °C at 222 nm. The reduction in the negative amplitude of the ellipticity indicates a reduced helical content (n=3, error bar=SD). The corresponding spectra are shown in Figure S2.

No significant changes in the proteińs fluorescence emission were observable until liposomes contained ≤40% PG. Beyond this threshold, the fluorescence intensity decreased progressively with increasing PG content, reaching a plateau at ∼70% PG (Figure 2C). This indicates that membrane binding of IM30 wt requires >40% PG to be monitored via fluorescence spectroscopy. Notably, the fluorescence emission characteristics of the wt protein changed only to a small extent upon membrane binding (Figure 2C).

IM30 wt forms large barrels in solution (Figure 1E, F), which disassemble upon binding to a negatively charged solid-supported membrane, as recently shown via AFM (Junglas et al., 2020). However, barrel formation is not a prerequisite for membrane binding, and IM30*, a variant defective in barrel assembly, also binds effectively to negatively charged membranes (Heidrich et al., 2016; Junglas et al., 2020). In fact, the fluorescence changes were more pronounced in case of IM30* compared to the wt, when the protein was added to PC/PG liposomes (Figure 2B, C). While fluorescence alterations were negligible below 40% PG for both variants, IM30* showed further binding when PG contents exceeded 70%, in contrast to the wt. The pronounced decrease of the IM30* fluorescence emission suggests that either the environment of the sole Trp in the membrane-bound state differs between IM30 wt and IM30*, or that solely a small fraction of the Trp’s contacts the membrane in the case of the wt protein. The latter explanation aligns with the wt’s stable barrel structure, where potentially solely the proximal barrel layer contacts the membrane, leaving most Trp residues unaffected (as further discussed below). Together, these observations confirm that barrel formation is not essential for membrane binding of IM30 and that the membrane-bound structures of IM30 wt and IM30* appear to differ to some extent.

Given the most pronounced effects were observed at 100% PG for IM30*, subsequent analyses focused on this lipid composition to maximize observable effects, acknowledging that this is non-physiological.

### The IM30 structure rearranges upon binding to PG membranes

The protein fluorescence characteristics of IM30 wt were less affected by PG liposomes compared to IM30*, suggesting distinct membrane interaction modes. To assess the respective membrane binding affinities, we analyzed the interaction of a constant amount of IM30(*) with increasing PG liposome concentrations (Figure 2D, E). Both variants exhibited a steep initial decrease in fluorescence emission with rising lipid concentrations (Figure 2D), indicating a high membrane-binding affinity. The fluorescence emission maximum of IM30 wt barely changed and plateaued already at about 100 µM PG, while it strongly decreased in the case of IM30*. At 300 µM PG, where binding of IM30* plateaued, the fluorescence intensity at 335 nm was decreased by ∼20% for the wt protein and >60% for IM30*.

These results suggest near-complete membrane binding for both variants at [PG] >250 µM. However, the differences in the fluorescence emission intensity underscore structural divergence in their membrane-bound states. Given that IM30 wt forms barrels in solution, while IM30* exists as monomers or small oligomers (Figure 1E-G), we next aimed at comparing the membrane binding of IM30 and IM30* under conditions where both variants are monomeric or present as small oligomers.

IM30 wt barrels were recently disassembled using urea, with the oligomer stability monitored via light scattering (Quarta et al., 2024). Increasing the temperature also disrupts barrels, as evidenced by a sharp decline in light scattering at temperatures >40°C (Figure S3). At about 50 °C, the scattering signal has dropped to 50%, *i.e.,* the oligomeric assembly was largely destabilized. Thus, we next studied membrane binding of IM30 and IM30* at 50 °C, where the wt barrel is substantially destabilized (Figure 2D).

For IM30*, the fluorescence emission at 335 nm changed only marginally between room temperature (25 °C) and 50 °C upon addition of PG liposomes, with a slightly steeper fluorescence decrease at the elevated temperature, suggesting a somewhat enhanced membrane binding affinity (Figure 2D). In contrast, IM30 wt’s fluorescence characteristics at 50 °C more closely mirrored IM30*, implying that the IM30 oligomeric structure significantly influences membrane binding. This aligns with prior reports (Heidrich et al., 2016), again confirming that barrel disassembly alters IM30’s mode of interaction with membranes and that the mutations introduced in IM30* are not the main cause for the different fluorescence characteristics of the membrane bound state.

Next, we investigated secondary structure changes in both IM30 wt and IM30* upon membrane binding at 25 °C and 50 °C via CD spectroscopy (Figure 2E). In the absence of lipids, IM30* exhibited a higher 208/220 nm ratio compared to IM30 wt, indicating distinct secondary structures (Figure S2). For both proteins, increasing lipid concentrations initially reduced the CD signal amplitude at 222 nm, followed by a retrieval at higher concentrations, suggesting transient loss followed by partial recovery of α-helical structure (Figures S4, 2E). This trend was particularly pronounced for IM30 wt at 50 °C, aligning with the fluorescence emission data (Figure 2D, E).

Plotting the ellipticity at 222 nm against the PG concentration revealed temperature-dependent differences (Figure 2E). For IM30*, the binding curves were similar at 25 °C and 50°C up to 100 µM PG, with a steeper initial slope at 50 °C. At higher PG concentrations, a clear increase in ellipticity was observed at 50 °C, and a similar, yet weaker trend at 25 °C. IM30 wt displayed an analogous behavior, yet the minimum ellipticity (at around 100 µM PG) was much more pronounced at 50 °C. Both variants exhibited biphasic binding curves at 50 °C. This suggests that the conformational states of membrane-bound IM30(*) proteins depend on the density of membrane-bound monomers.

### Various membrane-bound states of IM30

To probe potential membrane-induced structural changes, we next monitored the ellipticity at 222 nm for both IM30 variants at increasing temperatures (Figure 3A, B). In the absence of lipids, IM30 wt and IM30* displayed sigmoidal denaturation curves with transition temperatures of ∼56 °C and ∼54 °C, respectively (Figure 3A, B). The similarity in transition points suggests that the CD signal of soluble IM30(*) mainly reports on the stability of the α1-3 helical-hairpin structure, consistent with prior reports (Quarta et al., 2024). In the presence of PG liposomes, IM30* lost the cooperative unfolding behavior (Figure 3B), indicating heterogeneous conformational states or independent domain unfolding. Notably, both variants retained partial secondary structure in the membrane-bound state even at high temperatures. At low temperatures, the denaturation curve obtained for the wt protein in the presence of PG liposomes was similar to that of the protein in the absence of membranes, albeit the α-helicity was slightly reduced. Yet, IM30 wt’s melting behavior changed abruptly at ∼55 °C, *i.e.* at around the proteińs transition temperature where the barrels disassemble (compare Figure S3), transitioning from sigmoidal (soluble protein) to linear (membrane-bound fraction) regimes. Thus, the membrane-bound state of IM30 wt behaves as IM30* when barrels disassemble.

**Figure 3:**
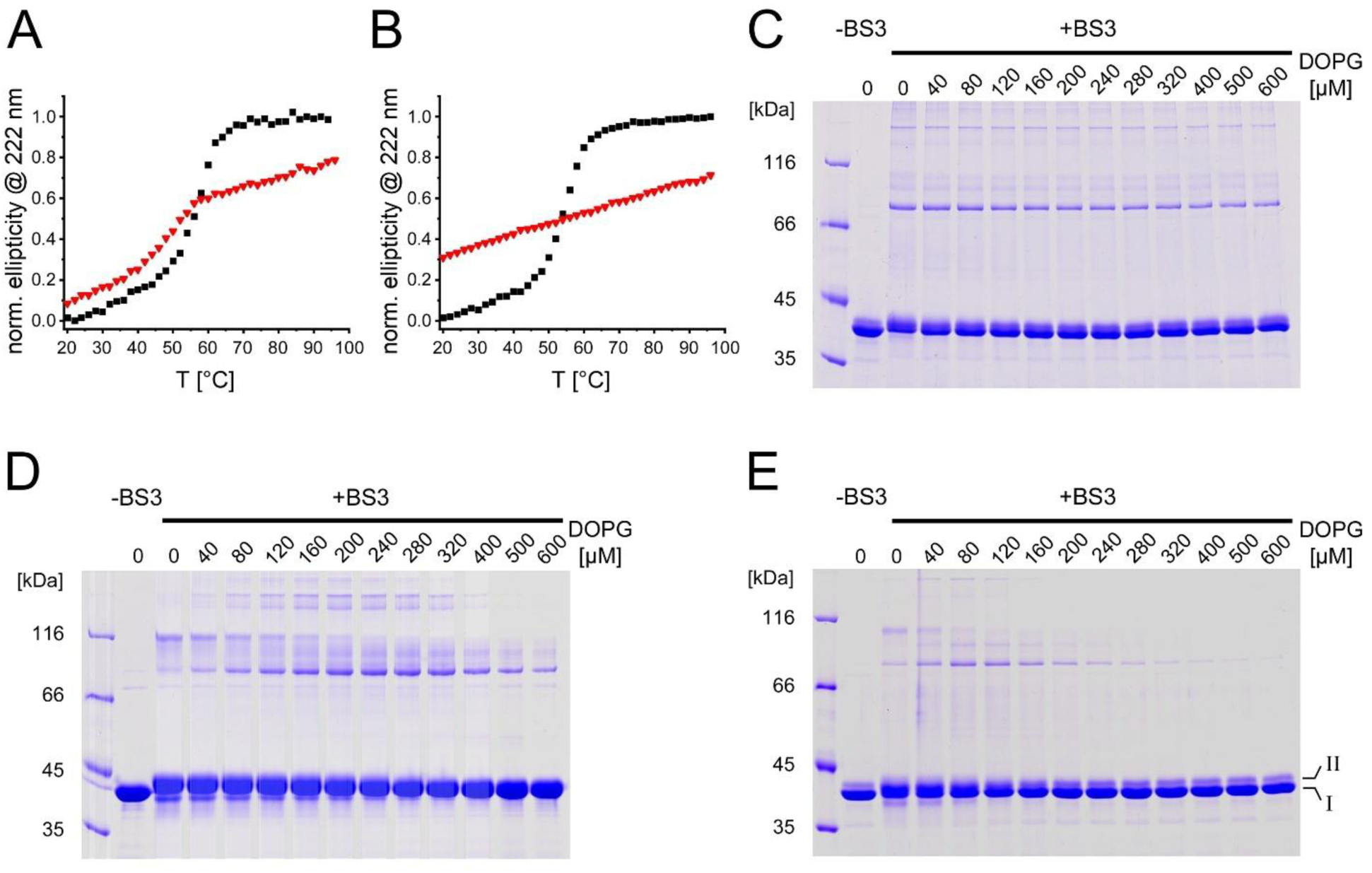
The secondary structure and oligomerization of membrane-bound IM30 are affected by elevated temperatures. (A, B) Ellipticity at 222 nm normalized to the value measured at 20 and 90 °C in absence of lipid during thermal denaturation of IM30 wt (A) and IM30* (B) free in solution (black squares) and in the presence of membranes (red triangles). For both proteins a classical sigmoidal curve is observed in solution. In the membrane-bound state, the curves of both proteins deviate significantly from this. (C-E) SDS-PAGE analysis of membrane-bound and subsequently cross-linked IM30 wt (C) and IM30* (D) at 25 °C, and wt at 50 °C (E). At 25 °C, IM30 wt shows monomer and oligomer bands that are independent of the protein density on the membrane. By contrast, at 50 °C the same protein displays a clear dependence on membrane protein density. The IM30* variant already exhibits this density-dependent oligomerization behavior at 25 °C. The two monomer species visible upon chemical cross-linking are labeled in (E). Calculated MWs: IM30 wt = 31.7 kDa; IM30* = 31.3 kDa.

To further monitor structural rearrangements upon membrane binding of IM30(*), we next aimed at cross-linking IM30(*) in the absence *vs.* presence of membranes to distinguish structural states during membrane binding (Figure 3C-E). For IM30 wt at 25 °C, BS3 cross-linking stabilized monomers and oligomers with apparent molecular masses of ∼40 kDa, ∼80 kDa, and ∼160 kDa, with no lipid-dependent changes, suggesting mainly preserved barrel structures (Figure 3C). At 50 °C, however, in IM30 wt samples two monomer species are visible: a dominant form (I) and a minor form (II) (Figure 3E). Increasing PG concentrations promoted oligomerization (∼80 kDa and ∼160 kDa), peaking at ∼80-120 µM PG before declining, indicating protein density-dependent interactions. A smaller oligomer of ∼116 kDa was observed at no or low PG concentrations.

At 25 °C, IM30* showed cross-linking patterns similar to IM30 wt at 50 °C, with prominent oligomers (116 kDa) at low PG concentrations (Figure 3D). Higher PG levels reduced the abundance of this oligomer and favored smaller (80 kDa) and larger (∼160 kDa) oligomeric species, as observed for the wt at 50 °C.

Both IM30 variants displayed (at least) two intramolecularly cross-linked monomer forms, with IM30*’s species II predominating at low PG and shifting to form I with an apparent lower mass at higher PG concentrations.

These findings demonstrate that membrane binding induces structural rearrangements in IM30(*), with conformational states depending on the protein density at the membrane surface. A high surface density promotes protein-protein interactions (visible as cross-linked oligomers). Even at high lipid concentrations, membrane-bound structures differed from solution states, confirming binding-induced conformational changes. The consistently observed biphasic binding behavior underscores differences in the structure of the bound protein at high and low surface density.

### The α1-3 helical hairpin is sufficient for membrane binding

Previously, it was proposed that membrane binding of IM30 is mediated exclusively by helix α0 (Ostermeier et al., 2024). However, our analyses of Trp fluorescence characteristics (Figures 2A-D) suggest that other helices are involved in IM30 membrane adhesion, including α1, which contains the only Trp residue (Figure 1A). Notably, α1-3 contains clusters of positively charged residues (Figure 1B), consistent with IM30 binding to negatively charged membranes. Given that helix α1, with its sole Trp residue, was not always present in subsequent analyses of IM30 fragments, we next employed an indirect approach to monitor membrane binding. This involved using membranes with the fluorescent dye Laurdan, which reports changes in membrane structure upon protein adhesion.

Comparing membrane binding of IM30 wt and IM30* revealed that the wt protein binds with lower affinity (Figure 4), consistent with previous measurements (Heidrich et al., 2016). Furthermore, the wt protein induced fewer alterations in membrane structure, as evidenced by a lower maximum GP value. This further supports the decisive influence of the IM30 oligomeric state on membrane binding.

**Figure 4:**
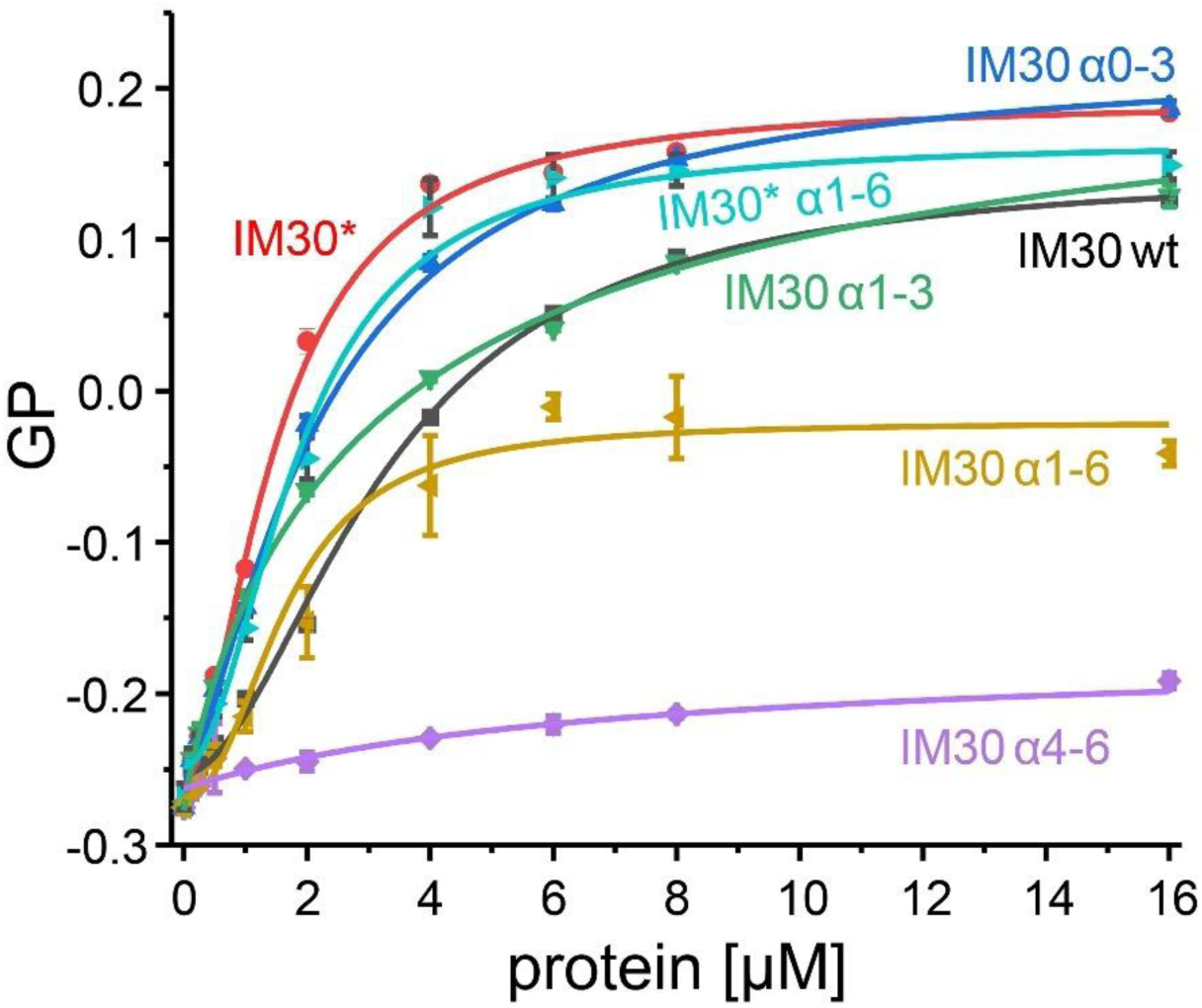
IM30 helices α1-3 mediate membrane binding of IM30. The Laurdan GP value at various concentrations of purified proteins (Figure 1D) was determined by fluorescence measurements. A change in GP value indicates a change in the microenvironment and thus the fluorescence emission of the Laurdan dye integrated in the DOPG liposomes. The GP derived binding curves were analyzed, and apparent KD values were determined: IM30 wt (KD = 3.27 ±0.17), IM30* (KD = 1.51 ± 0.12), IM30 α1-6 (KD = 1.65 ± 0.38), IM30* α1-6 (KD = 1.86 ± 0.17), IM30 α0-3 (KD = 2.20 ± 0.20), α1-3 (KD = 3.21 ± 0.84). For IM30 α4-6 no KD value could be determined (n=3, error bar=SD).

We then examined IM30 α1-6, where the N-terminal helix α0 was deleted. As shown in Figure 4, IM30 α1-6 still binds to membranes, albeit with a different impact on the GP value, *i.e.*, the membrane structure, compared to the full-length IM30 wt protein. Notably, while the wt protein primarily forms barrels in solution, IM30 α1-6 exclusively forms rods (Junglas et al., 2025; Thurotte and Schneider, 2019) (Figure 1E-G), which clearly influences its membrane binding properties. In fact, IM30* α1-6, *i.e.*, the variant not forming oligomeric structures anymore, bound to membranes essentially as well as full-length IM30* and with higher affinity than IM30 wt. These observations clearly confirm that α0 is not *per se* essential for IM30 membrane binding.

To pin down the helices involved in IM30 membrane adhesion, we next expressed and purified various truncated IM30 wt variants (Figure 1D, E). First, the full-length IM30 protein was split into two halves: IM30 α0-3, containing the structured α1-3 hairpin as well as the membrane-interacting helix α0, and IM30 α4-6, representing the disordered C-terminal part in solution (Junglas et al., 2020; Quarta et al., 2024). Additionally, we analyzed α1-3, the conserved helical hairpin structure without the N-terminal helix α0.

As presented in Figure 4, the addition of IM30 α4-6 to liposomes had only a small impact on the Laurdan GP value, indicating that the α4-6 region is not crucial for membrane binding. In contrast, the α0-3 variant bound with higher affinity to membranes than the wt, achieving GP values similar to IM30*, the monomeric IM30 variant with the exposed α1-3 hairpin (Heidrich et al., 2016; Junglas et al., 2020; Quarta et al., 2024). Furthermore, the α1-3 variant, lacking α0, still bound more effectively to membrane surfaces than the wt protein, although its membrane binding propensity was reduced compared to α0-3. Therefore, the α1-3 helical hairpin facilitates membrane binding of IM30, with α0 also contributing to this process, albeit to a minor extent, at least within the analyzed isolated system.

### The conformation of membrane bound IM30 a1-3 is governed by surface charge density

To further study the capability of α1-3 to bind negatively charged PG membranes in the absence of α0 and to elucidate the molecular mechanism governing this binding process, microsecond-scale all-atom molecular dynamics (MD) simulations of α1-3 in the presence of PC:PG membranes (at ratios 1:0, 1:1, 0:1) were performed. Spontaneous binding events occurred for all membrane types (Figure 5A), however, PG overall enhanced membrane binding of α1-3. First encounter events occurred earlier in membranes containing PG, resulting in an increase of the binding kinetic rate upon an increase in PG concentration (Figure 5B). Furthermore, binding occurred much more tightly and over prolonged periods with increasing PG content, as the number of protein–membrane contacts shifted to higher values when the concentration of this lipid was increased (Figure 5C). This confirms that α1-3 preferably associates with negatively charged membrane surfaces. Next, we analyzed which regions of the protein specifically interact with the membrane (Figure 5D). Consistent with the global protein–membrane contacts above, almost no sustained contacts were established between any residue and the pure PC (*i.e.*, 0% PG) membrane. However, as the PG content increased, not only were more binding events observed, but also the binding profiles changed: While at 50% PG concentration, mostly α1 was found to be in contact with the membrane, at 100% PG the binding shifted to a region closer to the N-terminal residues in α2/3. In both cases, mainly positively charged residues mediate the interaction.

**Figure 5.**
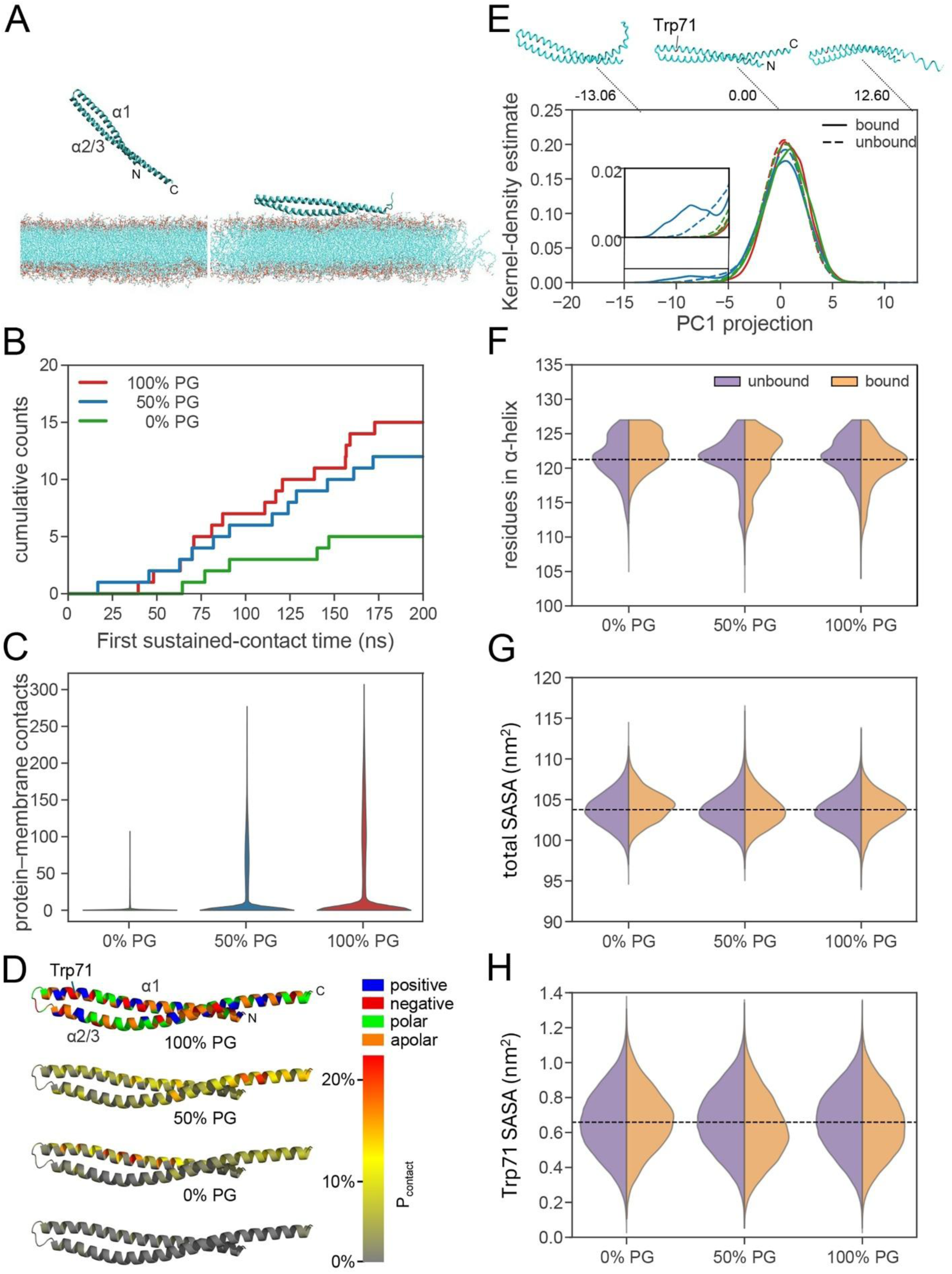
Association of IM30 α1-3 to PC:PG lipid bilayers was investigated by molecular dynamics (MD) simulations. (A) (Left) The α1-3 fragment (residues 26-156) was initially placed at a distant position of the PC:PG membrane (one example of such positions is shown here. (Right) Example of a spontaneous binding event observed during the MD simulations (in this case for a 1:0 PC:PG lipid ratio). Here, the protein is shown in cartoon representation, the lipid bilayer is depicted in stick representation. (B) Cumulative distribution function of the time elapsed to observe the first sustained-contact binding event for the indicated PG content in the lipid bilayer. Note that the maximum possible here is n=20, *i.e.* the number of replicas. (C) Distribution of protein–membrane heavy atom contacts for different amounts of DOPG in the lipid bilayer (violin representation). (D) Probability that a residue entered into contact with the membrane, Pcontact, is mapped on the color of the protein according to the color scale at the right (see Pcontact in 2D representation for each residue type in Figure S5). Amino acids are mapped above by type in colors. (E) Distribution of the projection of the simulation trajectories onto the principal component 1 (PC1). Projections are shown separately when the α1-3 fragment was either bound (solid lines) or unbound (dashed lines) to the membrane. Color represents the amount of PG in the membrane as in (B). Normalized KDE distributions are shown. The backbone configuration at two extreme projection values and at the center of the distribution is displayed above to visualize the collective motion represented by PC1. (F–H) Distribution of the number of residues in α-helical conformation (F), the total solvent accessible surface area (SASA) (G), and the SASA of residue Trp 71 (H) are displayed as a function of the PG content in the membrane, separately when the protein was bound or unbound to the membrane (violin representation). To guide the eye, the mean of the unbound fraction of the 0% PG (*i.e.* 100% PC) bilayer is marked with a dashed line.

In summary, our simulations support that the IM30 α1-3 membrane binding kinetics and affinity are enhanced in a PG concentration-dependent manner. However, the PG-to-PC ratio affects which residues preferably bind and thereby likely influences the membrane-bound protein conformation.

### An altered IM30 α1-3 conformation in the membrane-bound state

IM30 α1-3 membrane binding, and the structure of membrane-bound α1-3 were further evaluated using limited proteolysis, comparing fragment patterns in the absence *vs*. presence of membranes. In solution, without membranes, α1-3 proteolysis yielded several distinct cleavage intermediates, some of which are labeled in Figure 6A-C. Notably, the addition of PC liposomes to α1-3 resulted in an identical band pattern, reinforcing that IM30 does not bind to net uncharged membrane surfaces (Figure 6B). Species I, likely representing the monomeric full-length α1-3 fragment, over time transformed to species II, and both intermediates are persistent for a longer time in the absence of membranes or the presence of PC membranes (Figure 6A, B). Noteworthy, the transformation of species I to species II can be explained by the removal of C-terminal residues, as the N-terminal His_10_-tag was still present in the analyzed protein, as evidenced by a Western blot using an anti-His-tag antibody (Figure S6A-C). Only subsequently, digestions resulted in removal of the N-terminal His-tag, as any other species observed in the SDS PAGE analyses did not cross-react with the anti-His-tag antibody anymore (Figure S6A-C). Thus, the α1-3 C-terminus appears to be most sensitive to trypsin digestion, at least in the absence of liposomes and the presence of pure PC liposomes. The presence of PG liposomes significantly altered the band pattern, with species II becoming less prominent and shorter persistent (Figure 6C). Species I remained stable over an extended period before being degraded to species III, which also exhibited prolonged stability, a behavior not observed in the absence of membranes or when neutral (net-uncharged) PC membranes were present. This indicates that membrane interaction increases the overall proteolytic stability of α1-3, yet (at least) the α1-3 N- and C-termini were more accessible to the protease in the presence of PG liposomes, as the direct conversion to species III largely bypassed the intermediate form II. To characterize the proteolysis intermediate III that accumulated in the PG-containing sample, gel bands were cut, the protein fragment was hydrolyzed with trypsin, and the resulting peptide mixture was analyzed via liquid chromatography-coupled tandem mass spectrometry (LC-MS/MS). Database searching then confirmed the peptide sequences. Fragment III comprises approx. 57 amino acid residues (Leu77-Lys133), covering the loop 2 connecting helices α1 and α2, and essentially the entire helix α2.

**Figure 6:**
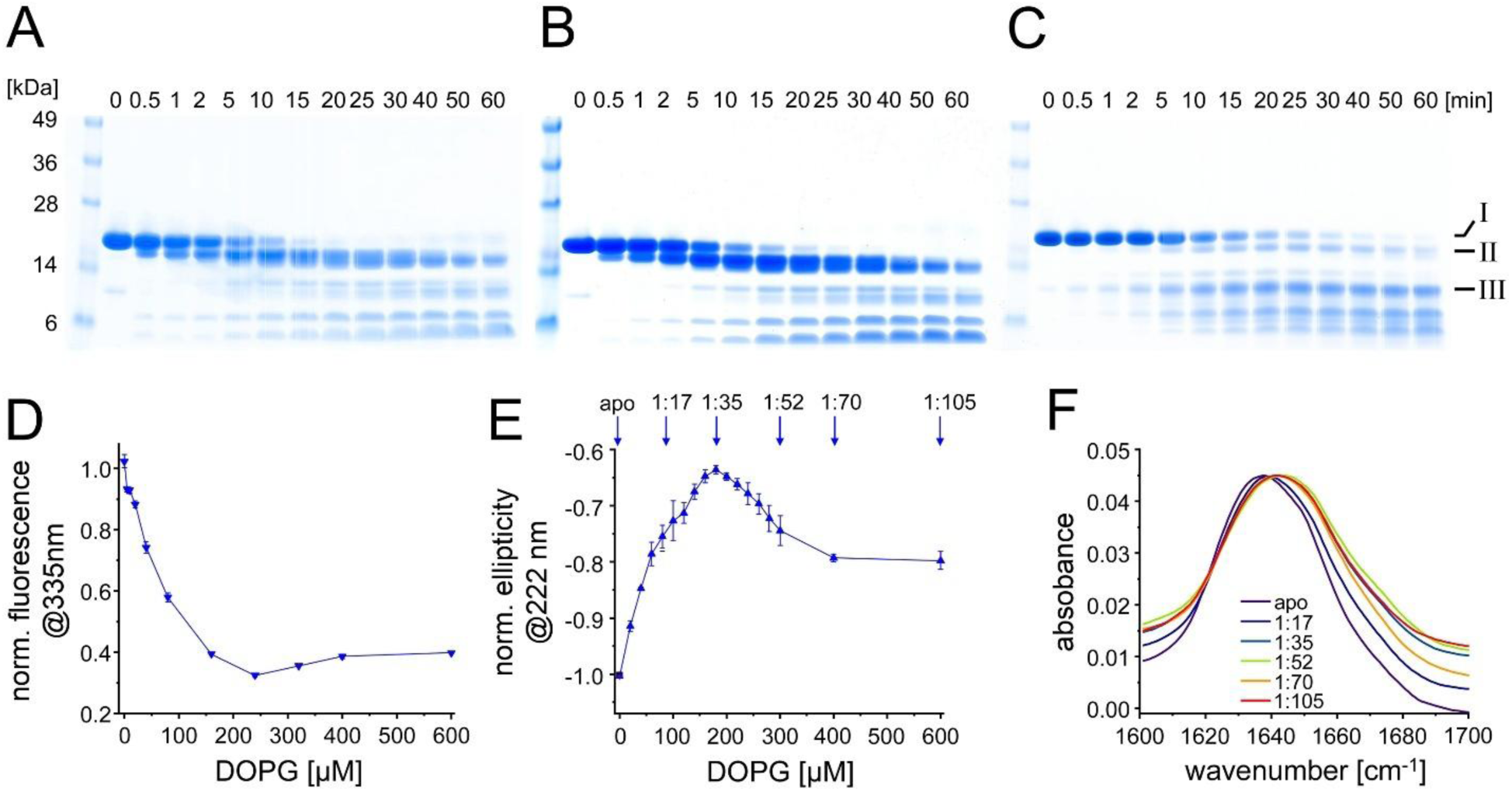
An altered structure of IM30 α1-3 in its membrane-bound state. (A-C) SDS-PAGE analysis of IM30 H1-3 in (A) absence of lipids or previously incubated with (B) PC or (C) PG liposomes followed by trypsin digestion. Samples were always taken before and 0.5, 1, 2, 5, 10, 15, 20, 25, 30, 40, 50, and 60 min after the addition of trypsin, as indicated in (A). I-V mark the proteolytic fragments of IM30 α1-3. (D) Normalized protein fluorescence intensities of IM30 α1-3 at 335 nm. A reduced fluorescence indicates membrane binding potentially coupled to rearrangement of the tertiary structure (n=3, error bar=SD). The full fluorescence spectra can be found in Figure S9. (E) Normalized ellipticity at 222 nm of IM30 α1-3 at increasing PG concentrations. A reduction in the signal intensity means a reduction in secondary structure content (n=3, error bars=SD). Blue arrows indicate the IM30 to PG ratios 1:17, 1:35, 1:52, 1:70 and 1:105 at each measuring point. The full CD spectra can be found in the appendix (Figure S9). (F) Normalized transmission FTIR spectra of IM30 α1-3 in solution and membrane-bound at certain protein-to-PG ratios. As the ratio decreases, the spectra shift to larger wavenumbers and become broader, indicating a decrease in α-helical structure content with increasing PG–to–α1-3 ratios until the entire protein is bound.

We further investigated to what extent the α1-3 conformation might change upon membrane binding by principal component analysis of the backbone motion calculated in our MD simulations (Figure 5E). The hairpin structure was conserved upon membrane binding and was rather independent of the PG fraction (see main peak of the distributions around zero of the projection in Figure 5E). However, when the fragment was bound to the 50% PG-containing membrane, it also adopted a second conformation in which the helical hairpin structure bent (see the second smaller peak at a value of approx. -8 in the distribution for this case in Figure 5E). In addition, the number of residues in an α-helical conformation decreased when the protein was bound to PG-containing membranes. As shown in Figure 5F, these bound states display a higher population in the distributions for values with lower α-helical content compared to their unbound counterparts and states bound to the 100% PC (0% PG) membrane. This reduction suggests partial unfolding upon binding. In fact, close inspection of the secondary structure of each residue over each simulation replica showed that these changes mainly occurred in the C-terminal region of α1-3 (Figures S6, 7). These changes, however, did not influence the global solvent accessible surface area (SASA) of the protein (Figure 5G), underlaying that the global structure of the α1-3 fragment was largely maintained upon binding. Nevertheless, locally, the SASA of Trp 71 decreased in the bound states compared to unbound states for PG-containing membranes (Figure 5H). The observed reduction in protein fluorescence in our experiments also indicates alterations in the local environment of this Trp residues (Figures 2, 3). Interestingly, all these effects were more pronounced for the 50% PG-containing membrane compared to 100% PG (compare 50% and 100% PG cases in Figures 5E, F, H). In summary, binding of α1-3 to PG-containing membranes bends and locally exposes part of the hairpin structure and partially unfolds the C-terminus of the α2/3 helix.

### Membrane-dependent conformations of IM30 α1-3

To experimentally investigate the indicated changes in the IM30 α1-3 structure upon membrane binding in more detail, we next examined Trp fluorescence emission changes of IM30 α1-3 at varying PG liposome concentrations (Figure 6D). The decrease in the Trp fluorescence emission intensity at increasing lipid concentrations was comparable between the α1-3 fragment and IM30* (compare Figure 2D), although the α1-3 fragment showed a slightly steeper decrease at low PG concentrations and a more pronounced increase above 240 µM. The similar membrane binding affinity of α1-3 and IM30* aligns with the previous observations: (i) IM30* does not form large oligomers (Figure 1E), (ii) a considerable disordered region (α4-6) in IM30* is not involved in membrane binding (Figure 4), and (iii) the α1-3 helical hairpin is stably structured and exposed in IM30* (Quarta et al., 2024).

We then investigated structural rearrangements associated with α1-3 membrane adhesion at the secondary structure level using CD spectroscopy (Figure 6E). The spectra, measured at increasing PG concentrations, indicated a biphasic binding behavior. Plotting the ellipticity at 222 nm against lipid concentrations revealed a minimal α-helical structure content at approximately 180 µM PG, with a subsequent increase in α-helicity at higher PG concentrations. To further elucidate these structural changes, transmission FTIR was employed (Figure 6F). Here, a much higher concentration of protein was required, yet, to be able to compare the results with the CD measurements, the protein/lipid ratios were maintained.

The FTIR spectra exhibited a shift to lower wavenumbers and broadened upon membrane binding, which became more pronounced with increasing lipid concentrations (Figure 6F). Above a 1:35 protein/lipid ratio, the spectra remained relatively constant, indicating a decrease in α-helical structure content with increasing PG–to–α1-3 ratios until the entire protein was bound. Notably, the slight increase in ellipticity observed in the CD measurements at higher lipid/protein ratios (Figure 6E) probably does not correspond to a regain of α-helical structure content, as the FTIR spectra did not show a corresponding backshift in wavenumber.

Via cross-linking, we further elucidated the structure of the α1-3 membrane-bound state (Figure 7A). Upon adding increasing amounts of liposomes to the isolated α1-3 peptide, a new monomer band emerged between the two monomer bands initially observed when α1-3 was chemically cross-linked in solution, and this new band became dominant when the PG concentrations were further increased. Notably, this new monomer band migrated at the same position as the non-crosslinked monomer on SDS-PAGE gels, suggesting that the residues cross-linked at intermediate PG concentrations were either inaccessible to the cross-linker or no longer in close proximity at higher PG concentrations. In contrast, the abundance of α1-3 monomers in the two states crosslinked in the absence of liposomes or at low lipid concentrations gradually decreased, with the lower species being almost entirely absent at high PG concentrations. While the Trp fluorescence measurements indicated that essentially all α1-3 was membrane-bound at lipid concentrations above 200 µM (Figure 6D), the cross-linking analysis now additionally revealed a change in the monomer structure upon membrane binding, as the band intensity of the non-crosslinked monomer species increased steadily with increasing lipid concentrations without reaching a maximum at approximately 200 µM lipid.

**Figure 7:**
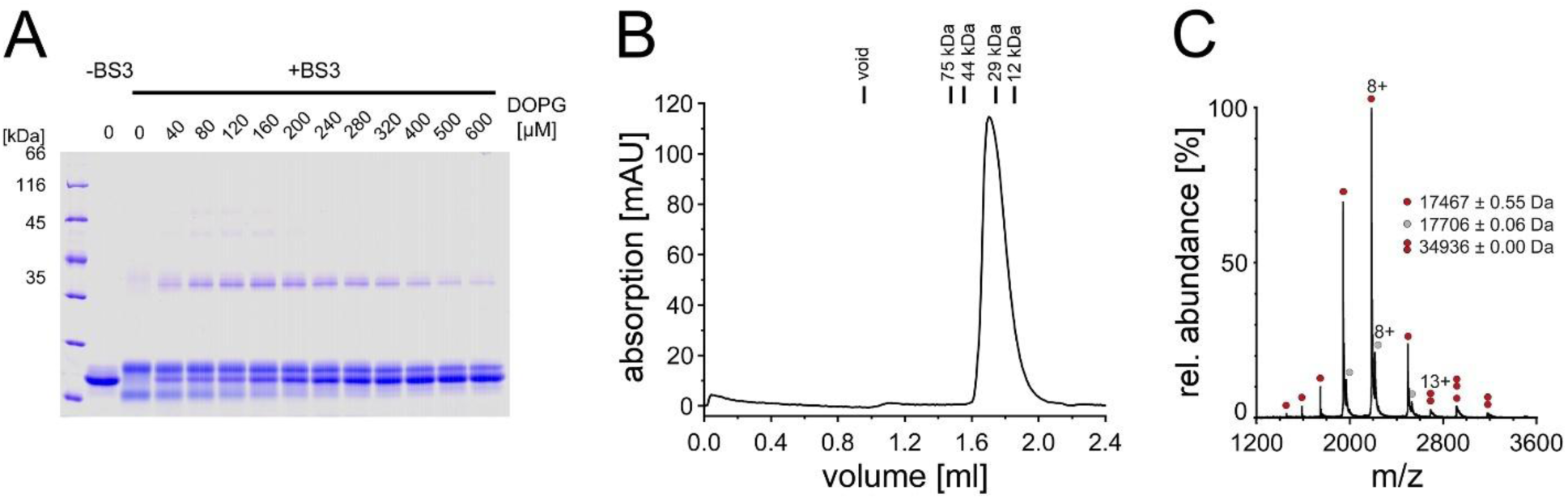
IM30 α1-3 adopts different conformations depending on the protein density on the membrane. (A) SDS-PAGE analysis of IM30 α1-3 without BS^3^ and after incubation with PG liposomes of increasing concentrations and subsequent BS^3^ crosslinked. BS^3^ cross-links primary amines, which occur in the side chains of Lys, among others. IM30 α1-3 monomers have a calculated mass of ∼17.5 kDa, yet run slightly higher at about 20 kDa in the absence of a crosslinker. After membrane binding, different monomeric IM30 α1-3 species, as well as higher ordered oligomeric species were cross-linked. (B) The chromatogram of an analytical gel filtration of IM30 α1-3 on a calibrated Superose 12 column shows a single dominant peak. (C) Native mass spectrometry showed that monomeric IM30 α1-3 is the most abundant species, with different charge numbers (single red dot). The second highest abundance is provided by IM30 H1-3 bound to a molecule of HEPES (gray dot). A further very small fraction of dimeric IM30 α1-3 was also detected (double red dot).

Furthermore, the dimer fraction exhibited a biphasic trend: as liposome concentrations increased, dimer intensity initially rose until approximately 160 µM PG, after which it declined with further increase in PG concentration. In addition to the dimer band, higher-ordered oligomeric species were observed at lipid concentrations between 40 µM and 200 µM. This suggests that at low lipid concentrations, where the liposomal surfaces were likely densely packed with α1-3, the proteins were in close proximity, promoting intermolecular crosslinking. The maximum dimer formation observed at approximately 160 µM lipid aligns well with the spectroscopically determined minima/maxima (Figure 6D, E), suggesting that the dimer is the species with the lowest fluorescence intensity and lowest α-helicity.

The low level of dimers observed in the absence of liposomes indicated that IM30 α1-3 is predominantly monomeric in solution. To investigate its initial oligomeric state in more detail, we conducted analytical gel filtration (Figure 7B). The chromatogram revealed a single dominant peak at 34 kDa, which could potentially represent a dimer based on the molecular mass of 17.5 kDa calculated for a monomer. However, since gel filtration chromatography requires a spherical protein structure for accurate size determination, and IM30 α1-3 has an extended structure deviating significantly from a sphere (Figure 1A, B), the results were inconclusive. To resolve this, we additionally employed native mass spectrometry, a technique that preserves non-covalent interactions in the gas-phase of the mass spectrometer and therefore allows determining the mass of non-covalently assembled proteins and protein complexes (Figure 7C). The mass spectrum showed one predominant charge state distribution corresponding in mass to monomeric IM30 α1-3. Adduct peaks of this charge state distribution revealed binding of a HEPES molecule (+238 Da). A second charge state distribution corresponding in mass to the dimeric protein was also observed albeit at very low intensity.

In summary, our combined experimental analyses indicate that (i) α1-3 is mainly monomeric in solution, (ii) upon membrane binding, α1-3 partially loses some secondary structure, accompanied by a decrease in protein fluorescence emission, and (iii) the extent of structural changes described in (ii) depends on the surface density of bound protein. The notable change in band patterns observed upon crosslinking the protein at varying protein/lipid ratios suggests that the distance and availability of cross-linkable residues are altered when α1-3 is membrane-bound. This alteration may stem from partial unfolding of α1-3, evidenced by CD spectroscopy (Figure 6E) and the simulations (Figure 5), or, alternatively, from partial destabilization of the coiled-coil structure upon membrane binding. The α1-2 hairpin structure is stabilized by interactions between hydrophobic residues located between α1 and α2, which may interact with the hydrophobic membrane core upon surface adhesion, potentially destabilizing the hairpin.

The ellipticity change and protein fluorescence both showed a biphasic behavior as the lipid concentration increased. At high lipid concentrations (i.e., low protein-to-lipid ratios) the spectroscopic signatures resembled those of the unbound protein, whereas at low lipid concentrations (high protein-to-lipid ratios) they differed markedly. Thus, the density of protein bound to the membrane surface influences the protein conformation.

At the concentrations used here, the surface density of bound protein was relatively high. Assuming a footprint for the α1-3 hairpin of ≈10 × 2 nm² and an area of ≈0.75 nm² per lipid head-group, a lipid-to-protein ratio of roughly 27 (≈1 protein : 54 lipids total) would be required to bind all protein. Achieving such coverage would demand an unrealistically ordered, side-by-side packing, which is entropically disfavored. For a rectangular molecule with a length-to-width ratio of 3, theoretical models predict a maximal coverage of only ∼20 % for random sequential adsorption (Minton, 1999).

If protein-protein contacts are established, however, the attainable surface coverage increases and can be reached at lower protein-to-lipid ratios. In general, higher surface densities shift the equilibrium toward conformations with a smaller footprint. The simulations suggest partial unfolding of the C-terminal residues, which would reduce the footprint and allow the termini to protrude from the membrane. This scenario is consistent with the CD data, although the simulated fraction of unfolded residues is smaller than observed experimentally, likely because the simulations lack interacting proteins.

In the FTIR experiments the protein concentration was about tenfold higher than in the CD measurements, pushing the equilibrium further toward the membrane-bound, densely packed state. Consequently, the helical signal does not re-appear at higher lipid concentrations (lower protein-to-lipid ratios) because the protein density remains sufficiently high.

Taken together, the data support a model in which membrane binding of α1-3 is mediated primarily by the α1-2 helical hairpin region, while the C-terminal helix tends to unfold. This unfolding enables a compact, ordered protein assembly that covers the membrane surface efficiently.

### The α1-3 helical hairpin mediates membrane binding of IM30 also in vivo

Thus far, membrane adhesion of the isolated α1-3 helical-hairpin has been exclusively shown and analyzed *in vitro*, in model membrane systems. To analyze, whether the isolated helix α1-3 hairpin is also sufficient for membrane adhesion *in vivo*, in living cyanobacterial cells, we next expressed wt IM30, IM30*, α0-3, α1-3 and α4-6, each fused to a yellow fluorescent protein (YFP), in living *Synechocystis* cells and monitored the subcellular localization via fluorescence microscopy. As can be seen in Figure 8, the wt protein was visible essentially exclusively as *puncta* structures, in line with recent observations (Bryan et al., 2014; Gutu et al., 2018). The IM30* protein was observed to be dispersed throughout the cytosol, with *puncta* formation also evident. Notably, a distinct localization of the protein at the cytoplasmic membrane became apparent, forming a halo-like structure around the cell. This observation can be readily attributed to the α1-3 hairpin exposed in the IM30* protein. In line with this, the α0-3 fragment behaved essentially as the IM30* protein, while the α1-3 fragment did not form *puncta* structures yet still localized to the cytoplasmic membrane. The α4-6 fragment remained uniformly distributed throughout the cytoplasm, as also observed for the free mVenus, supporting the low membrane binding affinity of α4-6 determined already *in vitro* (Figure 4). These findings align perfectly with and support our *in vitro* analyses, underscoring that the α1-3 helical hairpin is an important mediator of IM30 membrane binding, *in vitro* as well as *in vivo*.

**Figure 8:**
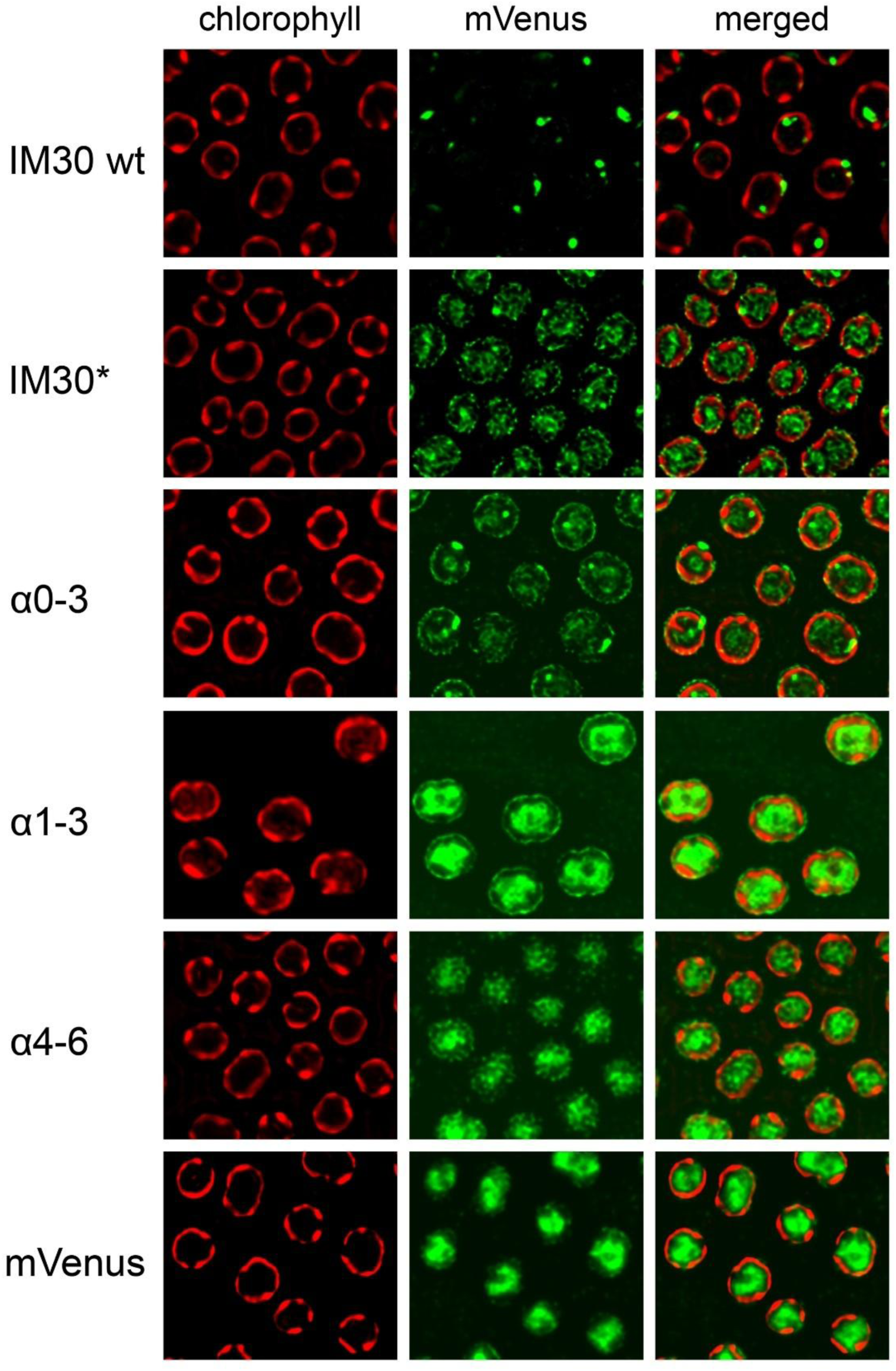
Membrane binding of IM30(*) variants *in vivo*. IM30 wt and variants were genetically fused at the respective C-terminus to a YFP separated by a 7GS linker. Subcellular localization and membrane binding were monitored via fluorescence microscopy. Chlorophyll fluorescence representing TMs is colored in red (1^st^ column) and mVenus fluorescence signals are colored in green (2^nd^ column). In the 3^rd^ column, the merged micrographs are shown. Scale bar = 2 μm.

## Discussion

In this study, we systematically investigated the binding of IM30 to negatively charged membrane surfaces. The helical hairpin formed by helices α1-3 is crucially involved in membrane binding, plus IM30 undergoes pronounced conformational changes upon binding to negatively charged membranes, affecting its secondary, tertiary, and quaternary structure. These structural rearrangements are influenced by the initial oligomeric state of the protein in solution as well as its surface density on the membrane.

### Membrane interaction of full-length IM30 depends on the oligomeric state and membrane surface coverage

IM30 has been demonstrated to bind to negatively charged membranes (Figures 2 and 4). Our *in vitro* analysis revealed that model membrane systems require a minimum of 40% negatively charged lipids to monitor IM30 membrane interactions through changes in protein fluorescence (Figure 2). Of note, with different techniques membrane binding of IM30 wt has already been observed at PG concentrations as low as 20% (Heidrich et al., 2016; Hennig et al., 2015). IM30*, a variant where oligomer formation is impaired (Heidrich et al., 2016; Junglas et al., 2020) (Figure 1E), displayed a different binding behavior as IM30 wt. When incubated with PG-containing liposomes, IM30* binding was observed via monitoring protein fluorescence changes at ≥40% PG (Figure 2C), followed by a steady decrease in protein fluorescence emission, suggesting a continuous increase in protein binding. This contrasts with the saturation observed for IM30 wt at ∼70% PG, despite identical protein concentrations. The primary difference between IM30 wt and IM30* is the formation of stable barrel and rod structures by the wt protein (Figure 1E-G), whereas IM30* exists as monomers and small oligomers in solution (Heidrich et al., 2016; Junglas et al., 2020; Thurotte and Schneider, 2019). The *in vitro* findings are consistent with our *in vivo* observations (Figure 8). Specifically, while only a small proportion of IM30 wt appears to associate with the inner membranes of cyanobacteria, IM30* was notably observed to bind to the cyanobacterial cytoplasmic membrane, as evidenced by a distinct halo surrounding the cells (Figure 8). This suggests that (i) oligomerization is not a prerequisite for IM30 to bind to membrane surfaces, and, in fact, (ii) oligomerization appears to hinder membrane binding, in line with recent observations (Heidrich et al., 2016). This is likely due to the shielding of membrane-interacting regions within the homo-oligomeric supercomplexes formed by the wt protein.

The amphipathic helix α0, which has been shown to interact with membrane surfaces (Junglas et al., 2025; McDonald et al., 2017), lines the inner surface of IM30 barrels and rods and is freely accessible only at one opening of the barrels (Gupta et al., 2021; Junglas et al., 2025; Liu et al., 2021). This suggests that membrane adhesion of IM30 rings may be mediated by helix α0. Although multiple α0 helices are present in the central lumen of IM30 barrels that have formed in solution, their involvement in membrane interactions is unlikely, as also observed in the homologous protein PspA (Hudina et al., 2025). PspA’s membrane internalization and tube formation involves initial disassembly of the oligomers, followed by membrane binding and oligomerization, which is coupled with membrane internalization (Hudina et al., 2025). Thus, the α0 helices within preformed IM30 barrels are probably largely inaccessible for membrane interaction as they are not in contact with a membrane surface. This explanation is consistent with the enhanced membrane binding affinity observed for IM30* (Figures 2, 4). As IM30* does not form stable barrel structures, all α0 helices are available for membrane binding. However, as shown in Figure 2D, E, the membrane interaction and structure of membrane-bound IM30(*) monomers depend on surface coverage. At high protein-to-lipid ratios (limited membrane surface availability), IM30(*) is less structured compared to its free state in solution or when sufficient membrane surface is provided. This difference may arise from varying interactions between IM30(*) monomers when in close proximity or the existence of distinct membrane-bound states, where more and/or different protein regions interact with the membrane when sufficient surface area is available. Indeed, the cross-linking data shown in Figure 3C-E support the assumption that monomer interactions on the membrane surface change with increasing surface availability, as the number of oligomeric species clearly depends on the protein-to-lipid ratio. Furthermore, when helix α0 would exclusively be responsible for membrane binding, IM30 α1-6, with helix α0 removed, should not bind to membrane surfaces anymore. The removal of helix α0 changes the architecture of the oligomeric assembly, with IM30 α1-6 forming extended rod-like structures instead of the barrel structures seen for the wt (Figure 1F, G). Despite this structural alteration, deleting the membrane-interacting helix α0 does not diminish the protein’s affinity for the membrane. It does, however, affect how the membrane properties are altered, which is reflected in the lower saturating GP value (Figure 4). Furthermore, IM30* α1-6, *i.e.*, the monomeric IM30 variant with helix α0 removed, binds essentially as well to membranes as the full-length IM30*, suggesting that membrane interaction of monomeric IM30, or low molecular weight oligomers, does not mainly depend on α0-membrane interactions. As IM30(*) membrane binding could be monitored via fluorescence changes (Figure 2), and the sole Trp located in helix α1 (Figure 1A, C), this helix and potentially the entire hairpin formed by helices α1-3 in IM30 appear to be involved in membrane binding. Thus, multiple membrane-bound states may (co)exist where membrane interaction is mediated by helix α0, helices α1-3, or both, depending on IM30’s oligomeric state.

### The helical hairpin formed by α1-3 is essential for membrane interaction

Fragmentation of full-length IM30 revealed that the helix α1-3 hairpin is sufficient for proper membrane binding, and a fragment including helix α0 (α0-3) bound with about the same affinity, despite the recent observation that the isolated helix α0 interacts well with negatively charged membrane surfaces (Junglas et al., 2025; McDonald et al., 2017). Our simulations supported α1-3’s membrane-binding propensity. The presence of negatively charged lipids crucially controlled both the association kinetics and the binding affinity of α1-3 to lipid bilayers (Figure 5B, C). Remarkably, the protein regions involved in binding (Figure 5D) and the subsequent structural changes observed in the protein upon binding, *i.e.* bending of the helical hairpin structure, partial unfolding of the α2/3 C-terminal region and reduced exposure of Trp71 (Figure 5E-H), depended in a non-monotonic fashion on the PG fraction: the effect was most pronounced at a PC:PG 50:50 ratio. Positively charged peptides tend to bind tightly to the head-group interface of pure PG membranes, due to strong and long-lived contacts between positive residues and negative PG groups. However, in 50:50 mixtures the α1-3 could have interacted with the hydrophobic parts of the PC lipids, too, resulting in different protein conformations, as readily observed in our simulations. Similar findings have previously been reported for amylin (Dignon et al., 2017). In contrast to α1-3, significant membrane interaction was not observed for helices α4-6 in our experiments, despite the recent implication of the C-terminal helix α6 in membrane binding (Hennig et al., 2017). It is possible that membrane adhesion of this helix requires the formation of supramolecular structures, which promote avidity. In line with the *in vitro* observations, we solely observed interaction of fluorescently labeled IM30 fragments with the cyanobacteria cytoplasmic membrane when the helix α1-3 hairpin was present and exposed, as in case of IM30*, IM30* α1-6, α0-3 and α1-3. Our combined analyses demonstrate that (i) the exposed helix α1-3 hairpin binds to membranes with high affinity, and (ii) helix α0 potentially further enhances this process.

The α1-3 helical hairpin has a calculated pI of 9.43, attributed to its excess content of Arg and Lys residues (Figure 1B). This composition explains the preferential interaction of the hairpin with negatively charged membrane surfaces. Spectroscopic (CD and Trp fluorescence, Figure 6D-F), proteolytic (Figure 6A-C) and computational (Figure 5) analyses revealed structural alterations in the hairpin upon membrane binding. The secondary structure and potentially tertiary contacts were clearly impacted by membrane interaction, as additionally evidenced by distinct fragment patterns observed upon incubation of α1-3 with trypsin in the presence of negatively charged membranes, compared to the absence of membranes or the presence of net-neutral PC membranes (Figure 6A-C). Although bound to a membrane surface, where shielding from protease might be expected, some cleavage sites became more accessible to trypsin than in solution. This strongly suggests that the protein is destabilized in regions containing these cleavage sites, again indicating partial unfolding of the α1-3 fragment upon membrane binding. Furthermore, a biphasic membrane binding behavior of α1-3 was observed. At an α1-3-to-lipid ratio of approximately 1:35, the high protein density led to the formation of higher-order, cross-linkable oligomers (Figure 7A). When the membrane was densely packed with α1-3, CD spectroscopy (Figure 6E) indicated a protein structure that is, compared to solution, more strongly altered than at lower packing densities.

Given that the helix α1-3 hairpin is stabilized by leucine-zipper-type hydrophobic interactions, it is plausible that the hydrophobic residues involved in forming and stabilizing this helical hairpin contribute to binding IM30 to the hydrophobic core regions of the membrane. This process likely occurs after initial membrane interactions mediated by positively charged amino acids, resulting in rearrangement of the membrane-bound α1-3 structure. Based on the proteolytic data (Figure 6A-C), membrane interaction primarily involves helix 2, which was most shielded when PG membranes were present. Structural rearrangements were also observed with the full-length protein (Figures 2, 3), and thus, can likely be attributed to the structural alterations in the α1-3 region. While this likely is the main effect in case of IM30*, for the IM30 wt the observed structural rearrangements may additionally involve changes in secondary structure upon barrel disassembly, as recently described (Junglas et al., 2020; Quarta et al., 2024).

In conclusion, the helix α1-3 hairpin exhibits a high propensity to bind to negatively charged membrane surfaces, with membrane binding destabilizing the α1-3 hairpin structure, likely involving partial unfolding. However, as shown via monitoring thermal unfolding (Figure 3A), once bound to the membrane, the secondary structure of the helical hairpin is highly stabilized by the membrane environment, indicating the formation of defined protein-lipid interactions.

### IM30 has at least two membrane-interacting regions

Eukaryotic members of the ESCRT-III superfamily are suggested to undergo a structural change prior to membrane binding, involving rearrangements in the α1-3 region (Bajorek et al., 2009; Tang et al., 2015). In its "closed" conformation, the yeast ESCRT-III protein Snf7 cannot bind membranes due to an auto-inhibited structure where α5 folds back against the α1-2 hairpin, disrupting the continuity of helix α3 with α2 (Lata et al., 2008). Opening of the ESCRT-III structures involves a large-scale conformational rearrangement, resulting in an overall elongated structure. This process includes the displacement of α5 from the α1-2 core domain (Henne et al., 2012; Lata et al., 2008; Tang et al., 2015) and the disruption of intramolecular interactions between the basic N-terminal and acidic C-terminal regions. Consequently, hydrophobic and electrostatic surfaces are exposed, facilitating protein interactions and liberating an extended cationic membrane-binding surface (Tang et al., 2015). In Snf7, specific Lys residues on α2 and α3 are crucial for membrane interaction (Tang et al., 2015). Initially located in distinct α-helices in the closed state, upon opening, these residues become arranged on a continuous, elongated, solvent-exposed surface, ideal for interacting with negatively charged membranes. In membrane-bound oligomers, the electrostatic membrane-binding regions of all Snf7 protomers face the same direction, forming a continuous, positively charged membrane-binding interface. Thus, for the eukaryotic ESCRT-III Snf7, membrane binding is coupled with oligomerization on membrane surfaces, driven by a structural rearrangement from a closed to an open conformation. The N-terminal membrane-anchoring α-helix α0 likely further stabilizes Snf7 on membranes (Buchkovich et al., 2013). Similarly, membrane interaction in the Asgard ESCRT-IIIB protein is mediated by two distinct regions: the N-terminal portion of helix α1, which bears some resemblance to α0 in IM30, and the α3-α4 connecting loop, characterized by a continuous stretch of exposed positive charges (Souza et al., 2025). Also, in human ESCRT-III protein CHMP3 monomers and oligomeric assemblies, an extended positively charged surface, formed by helices α1 and α2, is exposed that has been implicated in membrane binding (Muziol et al., 2006).

These observations nicely align with the findings described here. In IM30, two regions capable of membrane interactions have now been identified. The isolated IM30 helix α0 has previously been shown to bind to membrane surfaces (Junglas et al., 2025; McDonald et al., 2017), with interactions between helix α0 and the membrane surface being crucial for the formation and stabilization of tubulated membranes within IM30 (and PspA) rings and rods (Hudina et al., 2025; Junglas et al., 2025). However, consistent with the observation that eukaryotic ESCRT-IIIs in the closed conformation (with helix α0 in principle available for membrane interaction) remain soluble and unattached to membranes, the initial binding of IM30 monomers to membrane surfaces likely requires the positively charged surface of the α1-3 hairpin (Figure 1B), with helix α0 supporting membrane binding, as reinforced by our measurements (Figure 4). However, as *e.g.* shown in Figure 8, full-length IM30 wt primarily exhibits a cytoplasmic distribution *in vivo*, with some *puncta* formations. A significant membrane-bound fraction of IM30 wt was not apparent in our fluorescence microscopy, despite the presence of the α1-3 helical hairpin in IM30 wt. Yet, *e.g.* Figure 2 shows that α1-3 is not freely accessible when IM30 wt monomers assemble into oligomeric structures. Consequently, only a minor fraction of IM30 wt is membrane-associated, undetectable via fluorescence microscopy.

In summary, the following picture emerges that accounts for the observed membrane interactions and subsequent structural assembly: Upon initial membrane binding of IM30 monomers, primarily mediated by α1-3, IM30 monomers laterally interact on the membrane surface, similar to eukaryotic ESCRT-IIIs, ultimately leading to the formation of spiral, barrel or rod structures on the membrane surface, as suggested recently (McCullough and Sundquist, 2025; Naskar et al., 2025; Pan et al., 2024). This process might be coupled with membrane internalization and the formation of tubulated membranes within these rods and barrels (Gupta et al., 2021; Junglas et al., 2025; Junglas et al., 2020; Junglas and Schneider, 2018; Liu et al., 2021). Thus, at least two membrane-interacting regions exist in IM30, and potentially in PspA, where the positively charged helical hairpin α1-3 mediates initial contacts of monomers or small oligomers with negatively charged membrane surfaces, initiating the formation of larger oligomeric assemblies. Upon the formation of barrel or rod structures on membrane surfaces, helix α0 is primarily responsible for interacting with tubulated membranes within these barrels or rods.

## Materials and Methods

### Cloning, expression and purification of IM30 variants

Cloning of plasmids for expression of wt IM30 and IM30* was described in detail recently (Fuhrmann et al., 2009a; Heidrich et al., 2016; Hennig et al., 2015; Junglas et al., 2020). Plasmids enabling expression of N- or C-terminally truncated IM30 variants were generated via Gibson assembly using the plasmid encoding *im30* wt as a template (Fuhrmann et al., 2009a). All IM30 variants were heterologously expressed in *E. coli* BL21 (DE3) with a His-tag, as described (Thurotte and Schneider, 2019). Upon protein expression, cells were harvested via centrifugation, resuspended in purification buffer (50 mM sodium phosphate, 300 mM sodium chloride, 20 mM imidazole, pH 7.6), and disrupted via sonication. Unbroken cells and cell debris were removed by centrifugation, and the supernatant was used for protein purification via Ni^2+^-affinity chromatography. Matrix-bound protein was washed with purification buffer containing increasing imidazole concentrations (20 mM, 50 mM, 100 mM), and the proteins were finally eluted from the column with buffer containing 500 mM imidazole. The buffer was exchanged to 20 mM HEPES (pH 7.6) using PD-10 columns (Cytiva, Munich, Germany), and subsequently, the proteins were concentrated using centrifugal filters (Merck, Darmstadt, Germany).

### Size exclusion chromatography

IM30 variants were analyzed via size exclusion chromatography using a Superose 12 10/300 column (GE Healthcare, Munich, Germany). 500 µl protein samples in HEPES buffer (20 mM, 100 mM NaCl, pH 7.6) were loaded onto the column and analyzed at 7 °C using an ÄKTA purifier 10 system (GE Healthcare, Munich, Germany) with a flow rate of 0.5 ml/min. For better comparability of the peaks in the chromatogram, each curve was normalized to the same area underneath. The apparent protein mass of IM30 α1-3 was determined via size exclusion chromatography in HEPES buffer (20 mM, 100 mM NaCl, pH 7.6) using a Superose 12 or a Superdex® 200 Increase 3.2/300 column (Sigma, Taufkirchen, Germany) and an ÄKTA purifier 10 system (GE Healthcare, Munich, Germany). 30 µl sample volume (16 µM protein) was injected onto the column at 7 °C with a flow rate of 0.03 ml/min. For protein size estimation, the following standards were used: blue dextran (>2,000 kDa), ferritin (440 kDa), β-amylase (200 kDa), alcohol dehydrogenase (150 kDa), conalbumin (75 kDa), bovine serum albumin (66 kDa), ovalbumin (43 kDa), carbonic anhydrase (29 kDa), ribonuclease A (14.7 kDa).

### Liposome preparation

The lipids DOPG (1,2-dioleoyl-sn-glycero-3-phosphoglycerol) and DOPC (1,2-dioleoyl-sn-glycero-3-phosphocholine), both purchased from Avanti Polar Lipids, Inc. (Birmingham, AL, USA), were dissolved in chloroform and mixed in the appropriate volumes, if necessary. The organic solvent was evaporated by a gentle stream of nitrogen gas, and any remaining traces of the solvent were removed by vacuum desiccation overnight. Subsequently, the generated lipid films were rehydrated in HEPES buffer to form liposomes. Large unilamellar liposomes were prepared by three freezing and thawing cycles followed by liposome extrusion through a filter with a pore size of 100 nm (Whatman plc, Buckinghamshire, UK).

### Protein fluorescence spectroscopy

IM30 variants (5.7 µM or 2 µM) were incubated with DOPG liposomes (varying concentrations, see the main text for details) in HEPES buffer for 2 h at 25 °C. If not mentioned otherwise, all fluorescence measurements were performed using a FluoroMax+ fluorimeter (Horiba Scientific, Kyoto, Japan) with an integration time of 0.1 s at 25 °C. Proteins were excited at 280 nm (slit width 1 nm) and fluorescence emission, which is dominated by Trp, was monitored from 300 to 450 nm (slit width 3 nm). The emission spectra were normalized to the mean intensity between 333 nm and 337 nm in the lipid-free (apo) spectrum.

### Fourier transform infrared (FTIR) spectroscopy

For FTIR measurements, DOPG liposomes were prepared in HEPES buffer as described, yet D_2_O was used instead of H_2_O to prevent any overlap of the bending mode absorption of H_2_O and the amide stretching absorption of the protein. Isolated IM30 H1-3 was lyophilized, redissolved in D_2_O/HEPES buffer to a concentration of 60 µM (?), mixed, and incubated with DOPG liposomes at different molar ratios (1:17, 1:35, 1:52, 1:70, 1:105) for 2 h at 25 °C. Samples were held between two 1-mm-thick CaF_2_ windows separated by a 50-mm Teflon spacer. The measurement was performed using a Vertex 70 FTIR spectrometer (Bruker Corporation, Billerica, USA) in transmission geometry. The spectra were recorded with a resolution of 1 cm^−1^ at frequencies ranging from 400 to 4000 cm^−1^. During the whole measurement, the sample compartment was purged with dry air.

### Circular dichroism spectroscopy

IM30 variants (5.7 µM) and DOPG liposomes (varying concentrations, see main text for details) were incubated for 2 h at 25 °C in 10 mM HEPES/NaOH buffer (pH 7.6). Circular dichroism spectra were recorded from 200 to 250 nm using a CD spectrometer J-1500 equipped with a MPTC-490S temperature-controlled cell holder (JASCO Corporation, Tokyo, Japan) with a bandwidth of 1 nm, a scan rate of 50 nm/min, and an integration time of 1 s at 25 °C. Five measurements were accumulated. The spectra were normalized to the ellipticity at 222 nm of the respective apo spectrum.

For monitoring thermal denaturation, protein samples were heated from 20 °C to 96 °C, with a temperature increase of 1 °C/min and a 30 s equilibrating time. The spectral range was 200 to 250 nm with a bandwidth of 1 nm, a scan rate of 100 nm/min, and an integration time of 1 s with 5 measured accumulations. For evaluation, the ellipticity at 222 nm was normalized between -1 and 0 at 20 and 96°C, respectively.

### Dynamic Light Scattering (DLS)

Large oligomeric protein structures, such as IM30 wt barrels in solution, were monitored via dynamic light scattering (DLS). IM30 variants (2 µM) were analyzed in HEPES buffer. Scattering of 300 nm light was detected using a spectrofluorometer FP-8500 (JASCO Corporation, Tokyo, Japan). For monitoring the stability of oligomeric complexes, the sample was heated from 20 to 95 °C in steps of 5 °C with a heating rate of 1 °C/min.

### Laurdan fluorescence spectroscopy

Laurdan (6-dodecanoyl-N,N-dimethyl-2-naphthylamine, Sigma, Taufkirchen, Germany) is a fluorescent dye that incorporates into a lipid bilayer. Changes in the microenvironment of Laurdan result in an altered fluorescence emission spectrum. During liposome preparation, Laurdan was mixed in a molar ratio of 1:500 with DOPG dissolved in chloroform.

IM30 variants (varying concentrations from 0.125 µM to 16 µM) and DOPG/Laurdan liposomes (100 µM) were incubated in HEPES buffer for 2 h at 25 °C. Laurdan was excited at 350 nm (slit width 1.5 nm), and fluorescence emission was monitored from 400 to 550 nm (slit width 3 nm). To further evaluate the spectral changes, the Generalized Polarization (GP) value was calculated from each spectrum using the fluorescence intensities at 440 nm (I_440_) and 490 nm (I_490_).

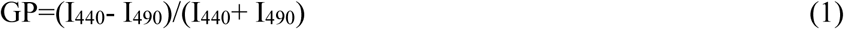

The apparent binding affinity (K_D_) was estimated by fitting the data with the following equation:

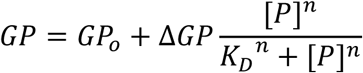

with [P] representing the protein concentration, GP_o_ the value in absence of protein, ΔGP the change in GP value upon protein binding.

### BS^3^ crosslinking

Prior to crosslinking, IM30 variants (5.7 µM) were incubated with DOPG liposomes at different lipid concentrations (40-600 mM) for 2 h at 25 °C. BS3 (bis(sulfosuccinimidyl)suberate, Thermo Fisher Scientific, Waltham, USA) was dissolved in HEPES buffer and added to a final concentration of 285 µM to the individual reactions, followed by incubation for 30 min at 25 °C. Subsequently, the crosslinking reaction was stopped with 50 mM Tris buffer (pH 7.6) for 15 min at 25 °C, and samples were analyzed via SDS-PAGE.

### Trypsin digestion

Prior to the trypsin digestion, IM30 α1-3 (5.7 µM) was incubated with DOPG or DOPC liposomes (final concentration: 200 µM each) and without liposomes for 2 h at 25 °C. Trypsin (Sigma, Taufkirchen, Germany) was dissolved in HEPES buffer. Both before the addition of 2.13 µg/ml trypsin and 0.5, 1, 2, 5, 10, 15, 20, 25, 30, 40, 50, and 60 min after the addition of trypsin, a sample each was taken from each preparation (with DOPG, with DOPC, or without liposomes) immediately mixed with 95 °C hot SDS-PAGE loading buffer to stop the digestion. Subsequently, the samples were analyzed using 4-12 % Bis-Tris gels of the NuPAGE system according to the manufacturers protocol (Thermo Fisher Scientific, Waltham, USA).

### Native mass spectrometry (MS)

Native MS experiments were performed on a Micromass Q-ToF Ultima mass spectrometer (Waters Corporation, Milford, USA) modified for transmission of high masses (MS Vision, Almere, Netherland) (Sobott et al., 2002). For this, the buffer of 30 µL protein solution was exchanged against 200 mM ammonium acetate using Micro Bio-Spin 6 gel filtration columns (BioRad Laboratories, München, Germany). The protein was then diluted to 10 µM, and 4 µL of the protein solution were loaded into gold-coated glass capillaries prepared in-house (Hernandez and Robinson, 2007) and directly introduced into the mass spectrometer. The following parameters were used for data acquisition: capillary voltage 1.7 kV, sample cone voltage 35 V, RF lens voltage 80 V, collision voltage 10V. Mass spectra were processed using MassLynx 4.1 (Waters Corporation, Milford, USA), externally calibrated using 100 mg/mL cesium iodide solution and analyzed using Massign software (version 11/14/2014) (Morgner and Robinson, 2012).

### Identification of peptides by LC-MS/MS analysis

Proteins were hydrolyzed in-gel as described previously (Shevchenko et al., 2006). Generated peptides were analyzed by reversed-phase liquid chromatography using a DionexUltiMate 3000 RSLC nano System coupled to a Q Exactive Plus Hybrid Quadrupol-Orbitrap mass spectrometer (Thermo Fisher Scientific, Waltham, USA). Mobile phase A was: 0.1% (v/v) formic acid (FA); mobile phase B was: 80% (v/v) acetonitrile (ACN), 0.1% (v/v) FA). Peptides were dissolved in 2 % (v/v) ACN, 0.1% (v/v) FA, loaded onto a trap column (Acclaim PepMap 100 C18-LC column, 300 μm I.D., particle size 5 μm; Thermo Fisher Scientific, Waltham, USA) and separated on an analytical column (Acclaim PepMap 100 C18-LC column, 75 μm I.D., particle size 3 μm; Thermo Fisher Scientific, Waltham, USA). For separation of the peptides, a gradient of 4-90% mobile phase B in 69 min was used. The following parameters were used for MS data acquisition: spray voltage 2.8 kV, capillary temperature 275 °C, data-dependent mode. Survey full scans were acquired in the Orbitrap (m/z 350-1600) with a resolution of 70.000 and an automatic gain control (AGC) target of 3e6. The 20 most intense ions with charge states of 2+ to 8+ were selected and fragmented in the HCD cell at an AGC target of 1e5 and a normalized collision energy of 30%. Previously selected ions were dynamically excluded for 30 s. The lock mass option (lock mass m/z 445.120025 76) was enabled (Hernandez and Robinson, 2007). For peptide identification, raw data were searched against a database including the protein sequence of IM30_H1-3 using MaxQuant (v.1.6.17) software (Cox and Mann, 2008). The following search parameters were applied: enzyme, trypsin/P; missed cleavage-sites, 2; fixed modification, carbamidomethylation (cysteine); variable modifications, oxidation (methionine) and acetylation (N-terminus); mass accuracy, 20 ppm for precursor ions and 4.5 ppm fragment ions; false discovery rate, 0.01.

### Negative stain electron microscopy

For negative staining electron microscopy (EM), 3.5 µl of the sample was applied to glow-discharged (PELCO easiGlow Glow Discharger; Ted Pella) continuous carbon grids (Carbon Support Film Cu-300 mesh; Quantifoil). The sample was incubated on the grid for 3 minutes. The grid was manually side-blotted using filter paper, washed with 3 µl of water, stained with 3 µl of 2% uranyl acetate for 30 s, and air-dried. The grids were imaged with a 120 kV Talos L120C electron microscope (Thermo Fisher Scientific/FEI) equipped with a CETA camera at a pixel size of 2.49 Å per pixel (×57,000 magnification) at a nominal defocus of 1.0–5.0 µm.

### In vivo localization and 3D rendering of fluorescently labeled IM30

For expression of mVenus-tagged IM30 variants in living *Synechocystis* cells, the IM30 constructs - featuring a 7GS linker preceding the mVenus tag at the protein’s C-terminus - were integrated into the plasmid pCK306. This plasmid enabled the insertion of chimeric genes into a non-essential region of the *Synechocystis* genome (the *ssl0410* locus) and enables a rhamnose-inducible expression of genes, driven by the *E. coli rhaBAD* promoter (Kelly et al., 2018).

*Synechocystis* wt cells were transformed with the pCK306-derived plasmids, and kanamycin-resistant clones were selected and cultured on BG11 agar plates containing increasing kanamycin concentrations. Successful segregation was verified via PCR analysis. The *Synechocystis* cultures were maintained in a shaker under continuous, low-intensity white light (30 μmol⋅photons m^−2^⋅s^−1^) in BG11 medium (Rippka et al., 1979), supplemented with 5 mM glucose and 100 µg/ml kanamycin. Prior to analyses, the cells were diluted to an optical density of 0.1 at 750 nm, and gene expression was induced by adding 1 mg/ml L-rhamnose. Following a 2-day incubation period (reaching an optical density of 0.4-0.6 at 750 nm), the cells were visualized using a ZEISS Elyra 7 microscope equipped with Lattice SIM2 super-resolution technology (Zeiss, Oberkochen, Germany), featuring a 63×/1.4 oil immersion objective. Imaging was performed using a two-channel setup, with a 642 nm laser for chlorophyll fluorescence and a 488 nm laser for mVenus excitation, with a filter set to SBS LP 560, allowing for simultaneous two-camera detection. Thirteen phase-contrast images were captured at 1280 x 1280 pixels with 16-bit depth. Super-resolution images were generated through SIM2 reconstruction through the Zen software. For 3D visualization, raw data was acquired at 0.110 μm intervals and processed using the SIM2 leap model, with 3D rendering performed in Imaris (version 10.1.1). Final image processing for this study involved adjusting saturation levels and incorporating scale bars using ImageJ (Rueden et al., 2017).

### Molecular Dynamics (MD) simulation parameters and algorithms

MD simulations were performed using GROMACS 2024.5 (Abraham et al., 2025) with the CHARMM36m (Huang et al., 2017) forcefield for the protein and lipids, the CHARMM TIP3P water model (Huang et al., 2017) and standard CHARMM ion parameters. A 1.2 nm non-bonded force cutoff with a Verlet cutoff scheme (Páll and Hess, 2013), and the LINCS constraint algorithm (Hess et al., 1997) for bonds involving the protein’s heavy atoms and the SETTLE algorithm (Miyamoto and Kollman, 1992) for bonds and angles of water molecules were used. Particle Mesh Ewald was used to compute electrostatic Coulomb forces (Darden et al., 1993). Simulations were carried out by maintaining both temperature and pressure constant, at values of 310 K and 1 bar, respectively, by using the v-rescale thermostat (Bussi et al., 2007) and the semiisotropic pressure C-rescale barostat. The time coupling constant of the thermostat was 1 ps and that of the barostat 5 ps. The reference compressibility for the pressure calculations was 4.5e-5 bar^-1^. Newton’s equations of motion were numerically integrated using the Leapfrog algorithm at discrete time steps of 2 fs (1 fs in the first equilibration steps of the systems containing membranes). Initial velocities were randomly generated from a Maxwell-Bolzman velocity distribution at 310 K.

### MD simulations of pure lipid bilayers

Three separate membrane bilayer systems with varying ratios of POPC and DOPG (1:0, 1:1, and 0:1) were produced using the CHARMM-GUI membrane builder (Feng et al., 2023), aiming for similar lateral dimensions after equilibration. To account for the compression of pure POPC bilayers and the expansion of pure DOPG bilayers during equilibration, systems were set up with lipid compositions of 686 molecules of POPC, 324 molecules of POPC and DOPG, or 600 molecules of DOPG, respectively. Systems were solvated with explicit water molecules, ensuring that a water layer of 13 nm surrounded the membrane (6.5 nm at each side). The membrane was neutralized by adding an excess of potassium ions. In addition, 0.15 M KCl was added to the water medium. Energy minimization, with steepest descent, followed by sequential equilibration steps in the NVT and NPT ensembles, at 310 K, was performed. During minimization and equilibration, position and dihedral restraints were imposed on selected atoms, and the strength of these restraints was gradually reduced as indicated in the CHARMM-GUI protocol. A final equilibration step of 100 ns, with all restraints lifted, followed. Final XY-dimensions of the simulation boxes oscillated around 14.5–15.5 nm. The final conformation was extracted as the initial coordinates of the lipid bilayer for later use.

### MD simulations of the IM30 α1–3 fragment in solution

The structure of α1-3 (aás 26 to 156) of the IM30 protein was extracted from the protein data bank (PDB id. entry 7o3y (Gupta et al., 2021)). The fragment was subsequently capped at the N- and C-terminus with neutral caps, thereby simulating its embedding in the full protein. Input parameters for the simulation were generated via the CHARMM-GUI solution builder (Brooks et al., 2009; Jo et al., 2008; Lee et al., 2016). A cubic simulation box was set up with at least a 1.5 nm buffer around the protein, with a total box length of 14.2 nm. The system was solvated using explicit water molecules, neutralized with an excess of ions. Additionally, 0.15 M KCl was added to the system. The protonation state of the titrable protein groups was chosen such that the pH was 7.0. During minimization, thermalization, and solvent equilibration, position restraints were imposed on backbone and sidechain atoms of 400 kJ/mol and 40 kJ/mol, respectively. Energy minimization with steepest descent to remove steric clashes was performed, followed by thermalization in the NVT ensemble at 310 K for 125 ps. Finally, position restraints on the protein were removed, and the dynamics of the protein were simulated for 200 ns in the NPT ensemble. From the resulting trajectory, 20 protein structures were extracted from equally spaced timepoints in the range between 40–200 ns.

### MD simulations of IM30 α1–3 in the presence of PC–PG lipid bilayers

Combined systems, considering the IM30 α1-3 fragment and the lipid bilayer, were generated using Python 3 with MDAnalysis 2.9.0 (Michaud-Agrawal et al., 2011)^15^. For each of the three lipid compositions, 20 simulation replicas were generated, each considering the protein in a different initial conformation and orientation relative to the lipid bilayer. In each replica, the equilibrated membrane without solvent was placed 1.5 nm above the bottom of the simulation box. The xy-dimensions were set equal to the inserted membrane simulation box, while the Z-dimension was set to 29 nm (sufficiently large to accommodate the protein in different orientations and the membrane). Subsequently, the protein was placed in the center of the box and rotated around its center of mass, ensuring the angle between the principal axis of the protein and the xy-plane did not exceed 45°. This was followed by a random rotation along the z-axis with a fixed random generator seed per replica across membranes. Lastly, the protein was translated along the z-axis such that at the end it was 2 nm above the membrane. Subsequently, both the protein and the lipid bilayer were explicitly solvated with water molecules, neutralized with an excess of potassium, and ionized with 0.12 M KCl. The number of atoms of the solvated systems varied from 515455 to 531668.

Possible atomic clashes were removed by energy minimization (5000 steps with steepest descent). Thermalization of the system was carried out in the NVT ensemble. Subsequently, the solvent and lipids were accommodated around the fragment in a series of MD steps (in the NPT ensemble), with position and dihedral restraints on lipids and the fragment. Restraints were gradually removed during such MD steps. The length and the strength of the geometric restraints at each successive equilibration step were chosen following the CHARMM-GUI protocol. 200-ns production runs followed, upon release of all position and dihedral restraints. 20 independent simulation replicas were considered, for a total of 4 microseconds of cumulative simulation time, for each lipid composition.

### Simulation analysis

Distributions of different observables of interest were generated by combining the data of the different simulation replicas. Analysis and visualization were performed using Python 3, GROMACS 2024.5, MDAnalysis 2.9.0 (Michaud-Agrawal et al., 2011) (Gowers et al., 2016), numpy (Harris et al., 2020), scipy (Jones et al., 2001), pandas (McKinney, 2010), seaborn (Waskom, 2021), matplotlib (Hunter, 2007), PyMOL 2.5.0 (Schrodinger, 2015), and vmd (Humphrey et al., 1996).

The minimum distance between the protein and the lipid bilayer and the number of contacts between the protein and the membrane were extracted from the simulation by the capped_distance function of MDAnalysis. A contact was recorded if any heavy atom of a protein residue came within a distance of 4 Angstroms of a heavy atom of the membrane.

The distribution of the number of contacts was computed separately for each lipid composition. In addition, the probability that each residue entered into contact with the membrane was estimated as the fractional occupancy per residue, *i.e.*, the total number of frames a respective residue presented at least one contact with the membrane divided by the total assessed simulation frames.

To assess the statistics, bootstrapping was performed with 1000 resamples, a confidence interval of 0.95, and the bias-corrected accelerated bootstrap interval for the 131 residues over 20 replicas using SciPy.

To estimate the kinetics of the binding process, binding was defined as >=5 heavy atom contacts between the protein and membrane maintained for >= 5 ns. This conservative criterion was chosen to avoid mislabeling transient contacts as binding events. As an indicator of binding time, the first time such sustained contact occurred was recorded for each simulation replica. These times were shown in a sorted and cumulative fashion. Unbinding was defined in a similar fashion, *i.e.* <5 heavy atom contacts between protein and membrane maintained for >=0.5 ns, and was assumed to only occur after a binding event. Via these binding and unbinding definitions, populations of the protein in each membrane system could be separated into bound and unbound states. Accordingly, the following structural properties of the protein were computed by distinguishing if the protein was in either of these two states.

Global structural changes of the protein backbone were assessed by principal component analysis (PCA). In brief, PCA consists of the calculation and diagonalization of the covariance matrix of the (here backbone) atomic positions (Amadei et al., 1993). The covariance matrix was computed and diagonalized via the GROMACS gmx covar function using the concatenated and fitted protein trajectories of the 1:1 PC:PG mixed membrane replicas. The C-terminal part of the fragment, namely residues 152 to 156, was found to be highly mobile and frequently varied its secondary structure (Figure S7). Thus, it was excluded from the PCA analysis, *i.e.*, only residues 26 to 151 were considered. The first principal component accounted for 58.97% of the total positional fluctuations. Subsequently, all three membrane-wise concatenated and fitted protein trajectories of each setup were projected along the eigenvector corresponding to this first principal component, via the GROMACS gmx anaeig function. Analysis of the secondary structure was performed using GROMACS gmx dssp function. The secondary structure for each residue was monitored as a function of time (Figure S8), and the number of residues that adopted a helical conformation at each time frame was computed.

Lastly, the solvent-accessible surface area (SASA) was computed for the whole protein and for the TRP 71 residue with GROMACS gmx SASA function, by rolling a probe solvent sphere of radius 0.14 nm on the protein surface.

Distributions were plotted with the seaborn package using the Scott method, either as Kernel density estimates (KDE) or in violin representation. For the violin plot of the number of residues adopting an alpha helical conformation, a smoothing factor of 2 was applied to the bandwidth.

GROMACS input parameters and scripts to analyze the trajectories can be found at the Github site https://github.com/graeter-group/IM30_alpha1-3.

## Supplementary material description

### Supplementary Figures S1-S9

**Figure S1:** Protein binding to PG liposomes monitored by changes in intrinsics protein fluorescence.

**Figure S2:** Membrane binding affects the secondary structure of IM30.

**Figure S3:** IM30 rings disassemble at elevated temperatures.

**Figure S4:** Membrane binding affects the thermal stability of IM30(*).

**Figure S5:** Fractional occupancy per residue of molecular dynamics simulations of IM30 α1-3 in presence of lipid bilayers

**Figure S6:** α1-3 is digested by trypsin from the N- and the C-terminus.

**Figure S7:** Secondary structure of each residue of IM30 α1-3 in presence of lipid bilayers recovered from MD simulations.

**Figure S8:** DSSP analysis of secondary structures of molecular dynamics simulations of IM30 α1-3 in presence of lipid bilayers (heatmap).

**Figure S9:** Membrane binding of IM30 α1-3.

## Author contributions

**Lukas Schlösser**: Conceptualization, Data Curation, Formal Analysis, Investigation (protein preparations and analyses), Methodology, Validation, Visualization, Writing – Original Draft, Writing – Review & Editing. **Mirka Kutzner**: Data Curation, Formal Analysis, Investigation (protein preparations and analyses), Validation, Visualization, Writing – Original Draft. **Nadja Hellmann**: Data Curation, Software, Validation, Visualization, Writing – Review & Editing. **Dennis Kiesewetter**: Conceptualization, Data Curation, Formal Analysis, Investigation (simulations), Methodology, Validation, Visualization, Writing – Original Draft. **Julia Bieber**: Conceptualization, Data Curation, Formal Analysis, Investigation (mass spectrometry), Methodology, Validation, Visualization, Writing – Original Draft, Writing – Review & Editing. **Ndjali Quarta**: Investigation (generation of fluorescently tagged strains). **Xingwu Ge**: Investigation (high resolution fluorescence microscopy measurements), Methodology, Visualization. **Tom Goetze**: Investigation (electron microscopy), Methodology, Writing – Original Draft, Writing – Review & Editing. **Benedikt Junglas**: Formal Analysis, Supervision, Writing – Original Draft. **Fumiki Matsumura**: Investigation (FTIR spectroscopy), Methodology. **Mischa Bonn**: Funding Acquisition, Project Administration, Methodology, Supervision, Writing – Review & Editing. **Frauke Gräter**: Funding Acquisition, Project Administration, Methodology, Supervision, Writing – Review & Editing. **Carsten Sachse**: Funding Acquisition, Project Administration, Supervision, Methodology, Writing – Review & Editing. **Lu-Ning Liu**: Funding Acquisition, Methodology, Project Administration, Supervision, Writing – Review & Editing. **Carla Schmidt**: Funding Acquisition, Methodology, Project Administration, Supervision, Writing – Review & Editing. **Camilo Aponte-Santamaría**: Formal Analysis, Supervision, Writing – Original Draft. Writing – Review & Editing. **Dirk Schneider**: Conceptualization, Formal Analysis, Methodology, Visualization, Funding Acquisition, Project Administration, Resources, Supervision, Writing – Original Draft, Writing – Review & Editing.

## Acknowledgments

This study was funded by the Deutsche Forschungsgemeinschaft (DFG, German Research Foundation, SA 1882/6-1, SCHN 690/16-1, and SFB1551 (Project number 464588647)). The project has received funding from the European Research Council (ERC) under the European Union’s Horizon 2020 research and innovation program (grant agreement No. 101002812) (to D.K., and F.G.) and from the Federal Ministry for Education and Research (BMBF, ZIK program, 03Z22HN22) (do C.S.). MD computations were performed on the HPC system Viper at the Max Planck Computing and Data Facility. Work in the lab of L.-N. L. was supported by the Biotechnology and Biological Sciences Research Council (BBSRC) (BB/Y01135X/1, BB/W001012/1), the Royal Society (URF\R\180030), and Leverhulme Trust (RPG-2021-286). We gratefully acknowledge the electron microscopy access time and computing time granted by the biological EM facility of the Ernst-Ruska Centre at Forschungszentrum Jülich. In this regard, we thank T. Heidler, S. Shazad, S. Singh, P. Sundermeyer and D. Baron for maintaining the electron microscopes.

## Conflict of interests

The authors declare no conflict of interest.

## Notes

### Competing Interest Statement

The authors have declared no competing interest.

