## supporting data figures S1-S9; Table S1 for "Membrane binding of a cyanobacterial ESCRT-III protein crucially involves the helix α1-3 hairpin conserved in all superfamily members"

[1]Department of Chemistry, Biochemistry, Johannes Gutenberg University, Mainz, Germany; [2]Max-Plack Institute for Polymer Research, Mainz, Germany; [3]Institute for Scientific Computing, Heidelberg University, Im Neuenheimer Feld 205, Heidelberg, 69120, Germany; [4]Institute of Systems, Molecular and Integrative Biology, University of Liverpool, Liverpool L69 7ZB, United Kingdom; [5]Ernst-Ruska Centre for Microscopy and Spectroscopy with Electrons, ER-C-3: Structural Biology, Forschungszentrum Jülich, Jülich, Germany; [6]Institute of Molecular Physiology, Johannes Gutenberg University Mainz, Mainz, Germany.

**Running title:** IM30/Vipp1 membrane interaction

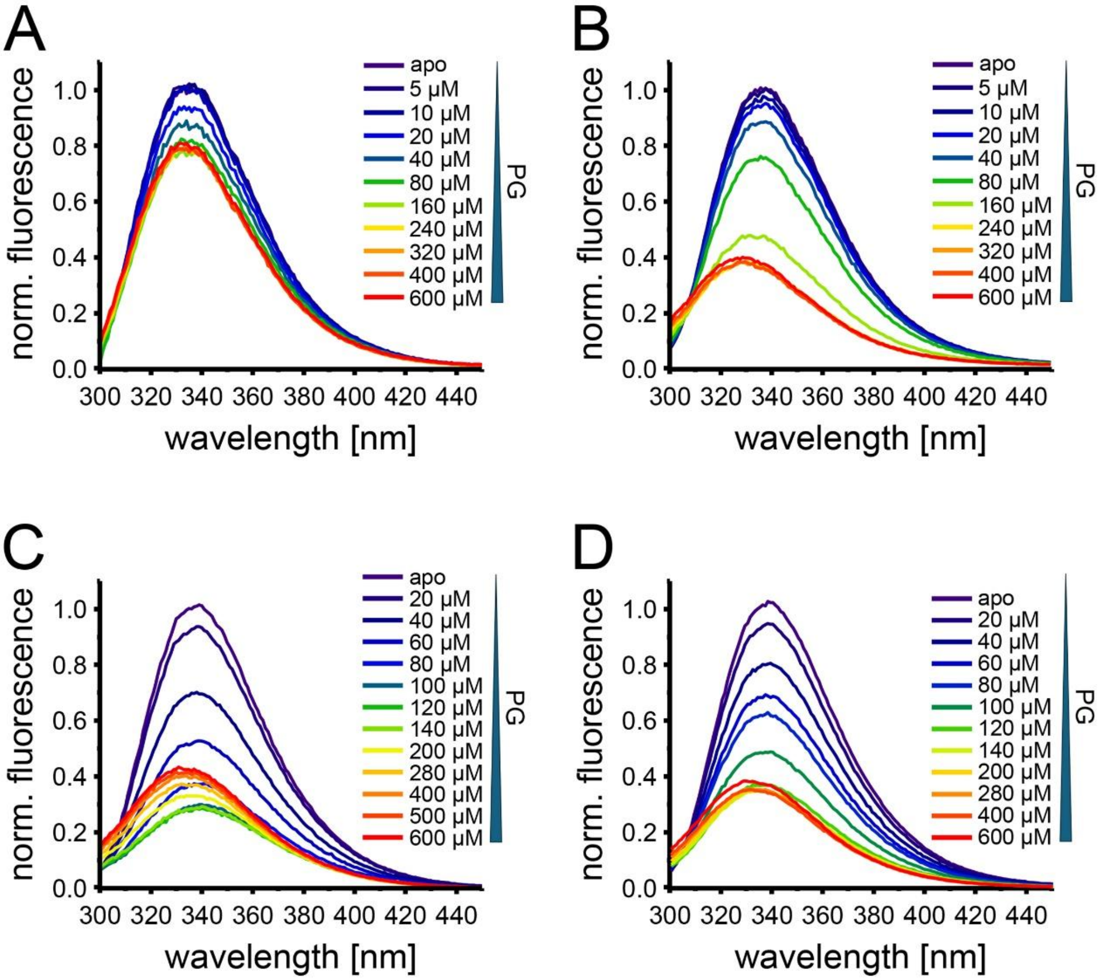


**Figure S1: Protein binding to PG liposomes monitored by changes in intrinsics protein fluorescence.**

Representative normalized spectra of the intrinsic protein fluorescence of IM30 wt (A, C) and IM30* (B, D) in solution (apo) and after incubation for 2 hours with increasing amounts of PG liposomes measured at 2 5°C (A, B) and 50 °C (C, D), respectively. Except in case of the wt at 25 °C (A), in all case a strong decrease in the fluorescence intensity is observed, accompanied by a blue-shift of the spectrum. This is mainly caused by the decreasing contribution of tryptophan fluorescence to the total emission. In particular in case of the wt protein measured at 50 °C (C), at a certain PG concentration the intensity increases again, and the blue shift increases, indicating that here clearly the environment of IM30's sole Trp has become more hydrophobic.

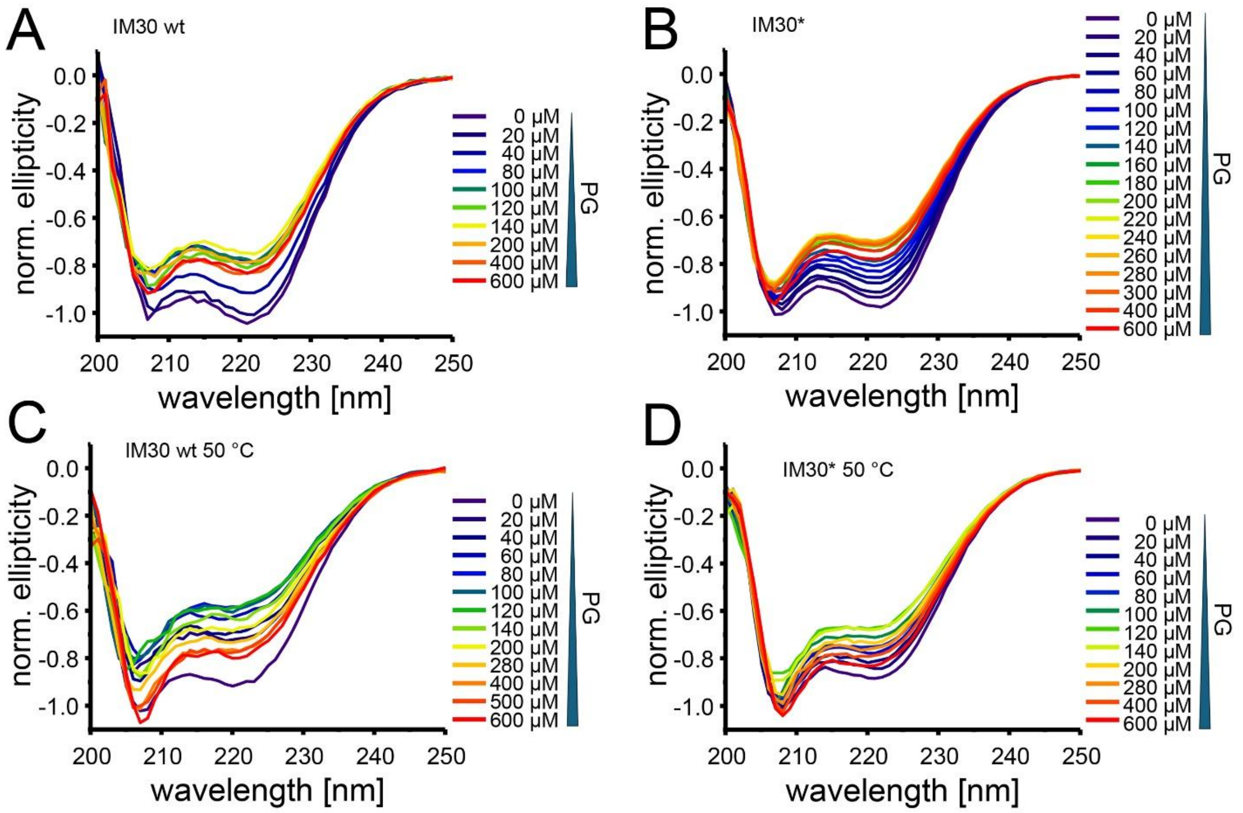


**Figure S2: Membrane binding affects the secondary structure of IM30.**

Representative normalized circular dichroism spectra of IM30 wt (A, C) and IM30* (B, D) in solution (apo) and after incubation for 2 hours with increasing amounts of PG liposomes measured at 25 °C (A, B) and 50 °C (C, D), respectively. In all cases the ellipticity at 208 nm and 220 nm initially decreases upon membrane binding, and increases again at higher PG concentration. Also, the shape of the spectra changes: the minimum at around 220 nm becomes shallower than the one at 208 nm in presence of PG liposomes. Overall, the effect on the spectra obtained for IM30* is more pronounced than for the wt protein.

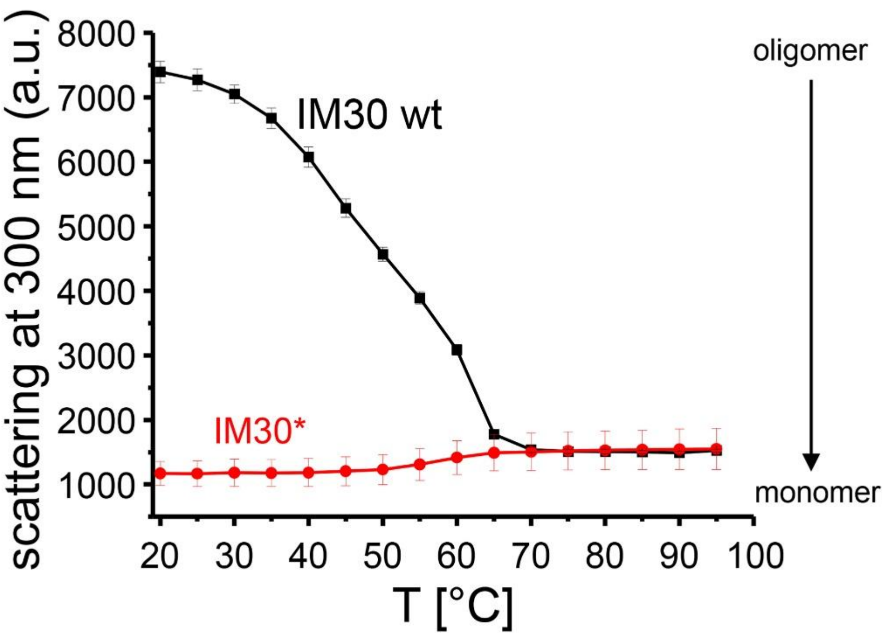


**Figure S3: IM30 rings disassemble at elevated temperatures.**

IM30 wt barrels were destabilized by increasing the temperature. The decrease of the scattering intensity clearly shows that the barrels disassemble already at relatively low temperatures. At about 65 °C the intensity is very similar to the scattering signal observed with IM30*, indicating the no large oligomers remain. However, as indicated by the slight increase of the scattering of the variant at this temperature, at temperatures >50 °C already denaturation and slight aggregation occurs. In order to stay clear of this temperature region, the experiments at elevated temperatures were performed at 50 °C. N=3, the error bars depict the SD.

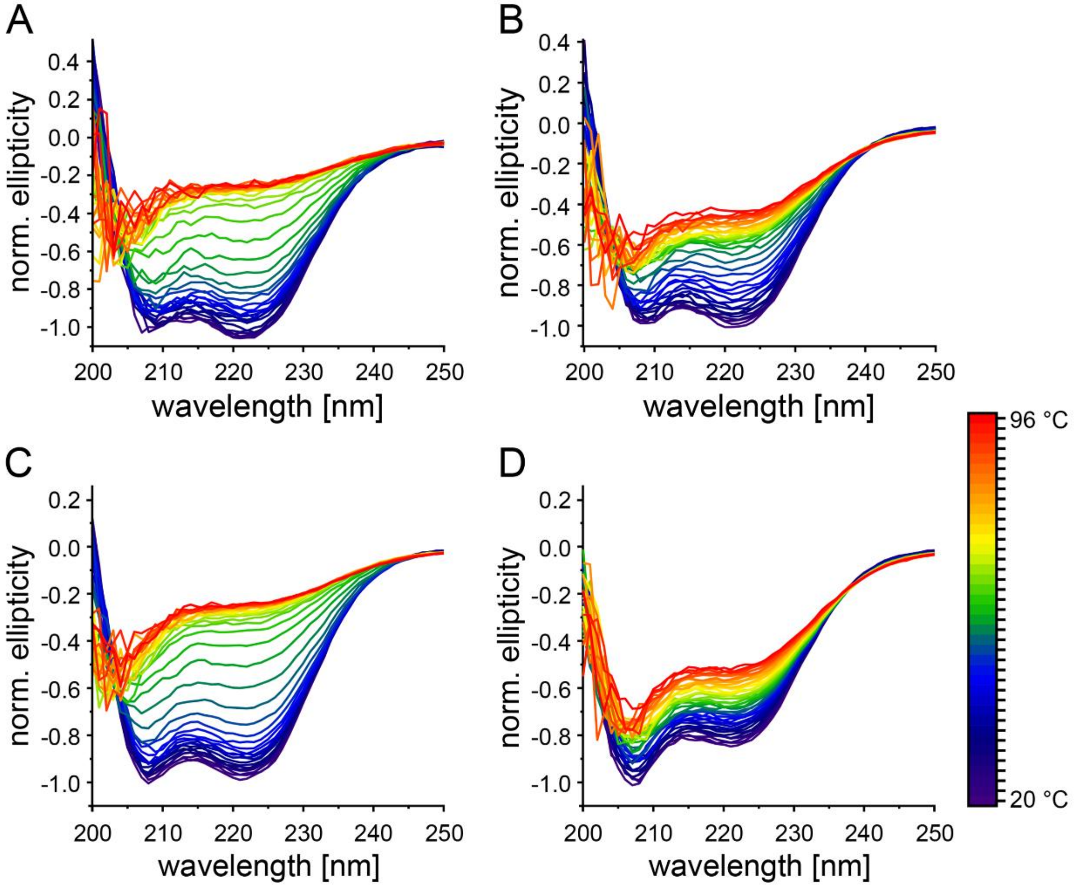


**Figure S4: Membrane binding affects the thermal stability of IM30(*).**

CD spectra of IM30 wt (A, C) and IM30* (B, D) at different temperatures, in absence (A, B) *vs*. presence (C, D) of 300 µM PG. The spectra were normalized at 220 nm. In absence of lipids, the transition between folded and unfolded protein is steep, while in presence of the lipid, it is much more gradual (compare Figure 3A, B). Also, complete loss of secondary structure is not achieved in the temperature range studied in presence of PG, as indicated by the residual signature of helical structures in the spectra.

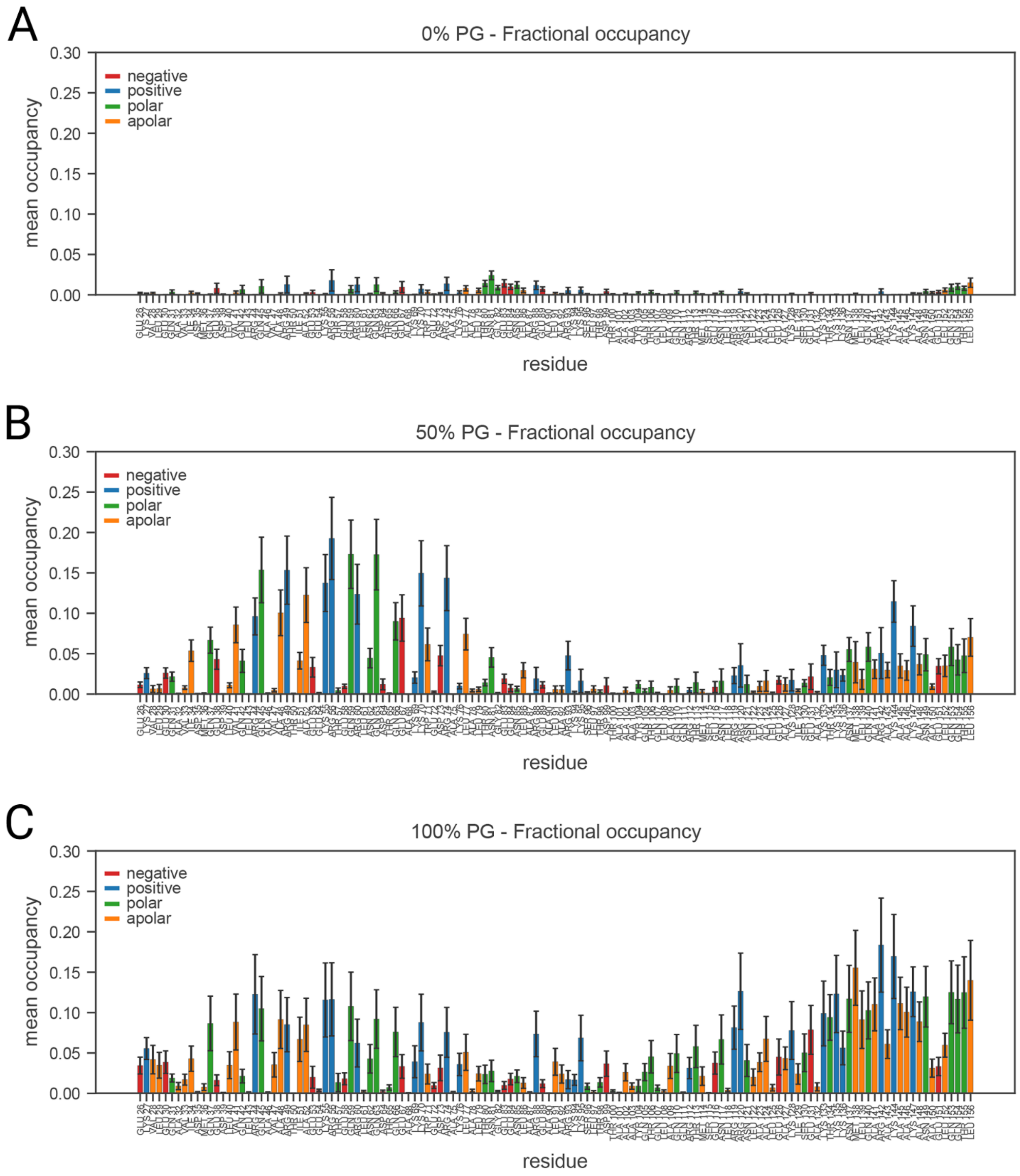


**Figure S5: Fractional occupancy per residue of molecular dynamics simulations of IM30 α1-3 in presence of lipid bilayers**

Statistical means and errors are computed via bootstrapping.

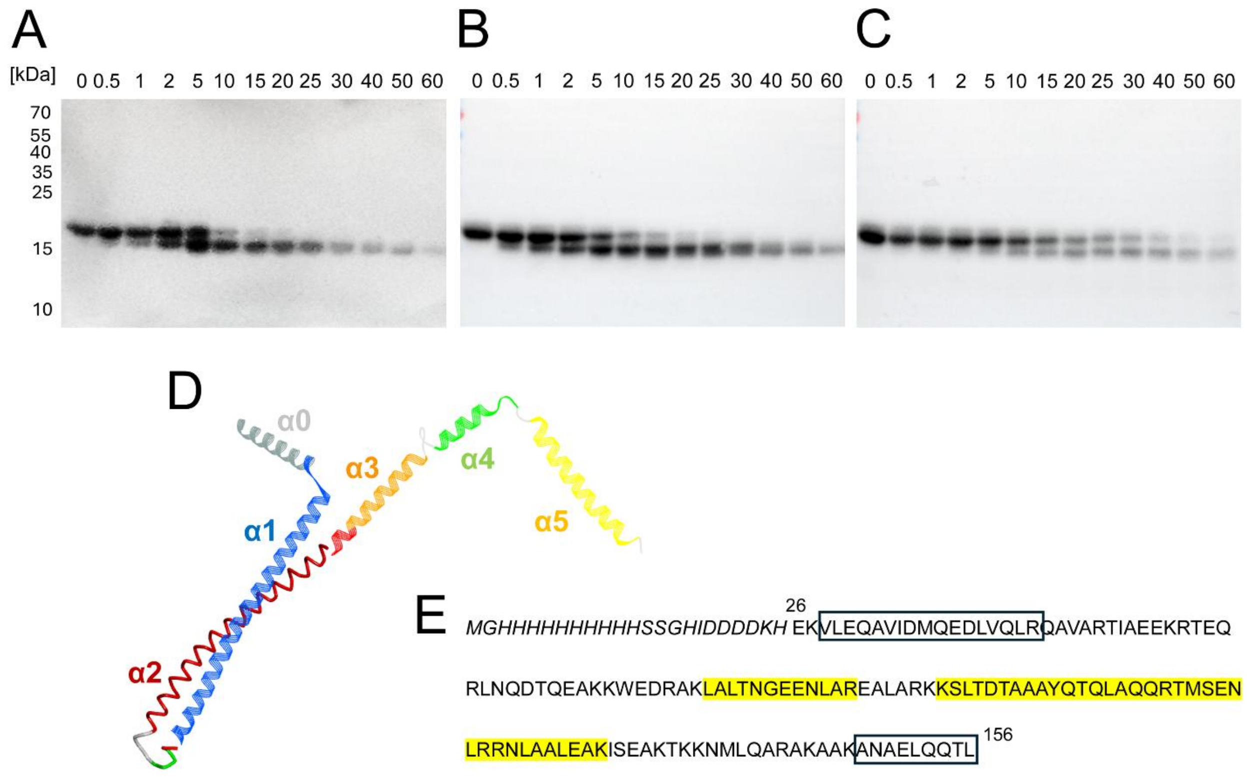


**Figure S6: α1-3 is digested by trypsin from the N- and the C-terminus.**

(A-C) Western blot analyzes of α1-3 upon trypsin digestion using an anti-His-tag antibody. The protein was digested (A) free in solution, (B) in the presence of PC liposomes, or (C) in the presence of PG liposomes. The full-length α1-3 with the N-terminal $His_{10}$-tag is first cleaved from the C-terminus, leaving the His-tag intact. The two fragments visible in (A) correspond to species I and II in Figure 6. Conversion of II to III involves removal of the N-terminal $His_{10}$-tag, as the anti-His-tag-antibody does not recognize fragment III any longer. (D) Structure (pdb: 7O3y) of IM30 wt with the helices colored as in Figure 1. Fragment IV protected upon membrane binding is highlighted in tube representation. (E) Sequence of the analyzed α1-3 fragment with an N-terminal $His_{10}$-tag plus linker (in italic). The analyzed IM30 fragment comprises residues 26 to 156. Sequences identified via MS analysis of the digestion intermediate IV, which was stabilized in the presence of PG membranes (compare Figure 6), are highlighted in yellow. For detailed MS analysis data see Table S1. Note that two peptides (boxed) were always identified in our analyzes, most likely as the physicochemical properties of this peptide favor ionization and fragmentation even at low abundance. Furthermore, due to high sensitivity of LC-MS/MS, overlapping species that could not be separated by gel electrophoresis will be identified even at low concentrations. The results of the Western blot analyses (A-C) clearly proof that α1-3 is initially cleaved from the N- as well as C-terminus.

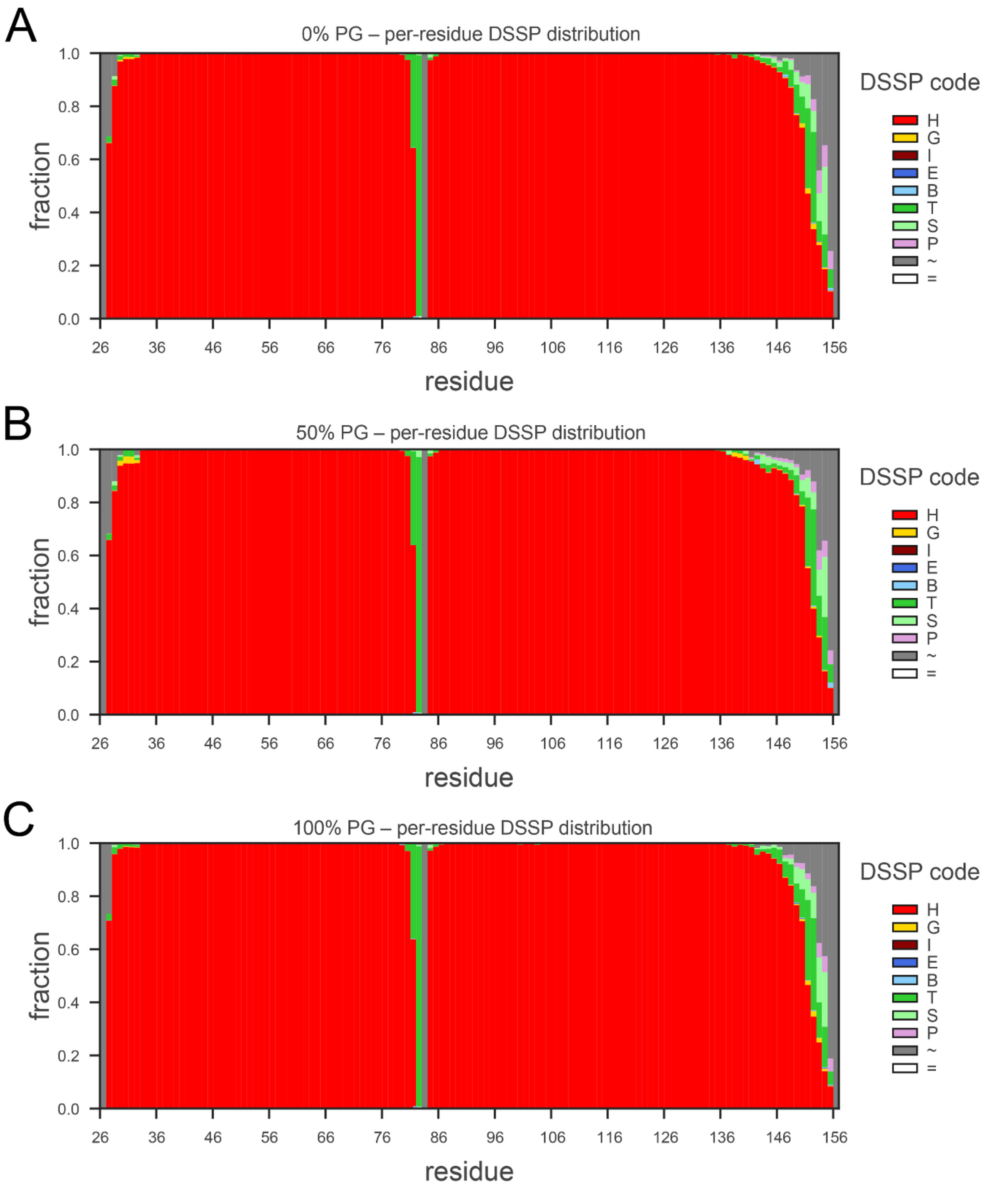


**Figure S7: Secondary structure of each residue of IM30 α1-3 in presence of lipid bilayers recovered from MD simulations.**

The x-axes indicate the residues of the peptide while the y-axes display the fraction of simulation time each residue adopted either of the indicated secondary structures. As visible in (A), (B) and (C) most of the peptide is conserved in a α-helical confirmation ("H", red) with the exception of the central loop connecting α1 and α2/3 with the C-terminal region in α2/3 frequently changes its confirmation indicating increased mobility and disorder compared to the rest of the peptide. Unfolding of alpha helical structures seems most pronounced in (B) in a 1:1 PC:PG mixture.

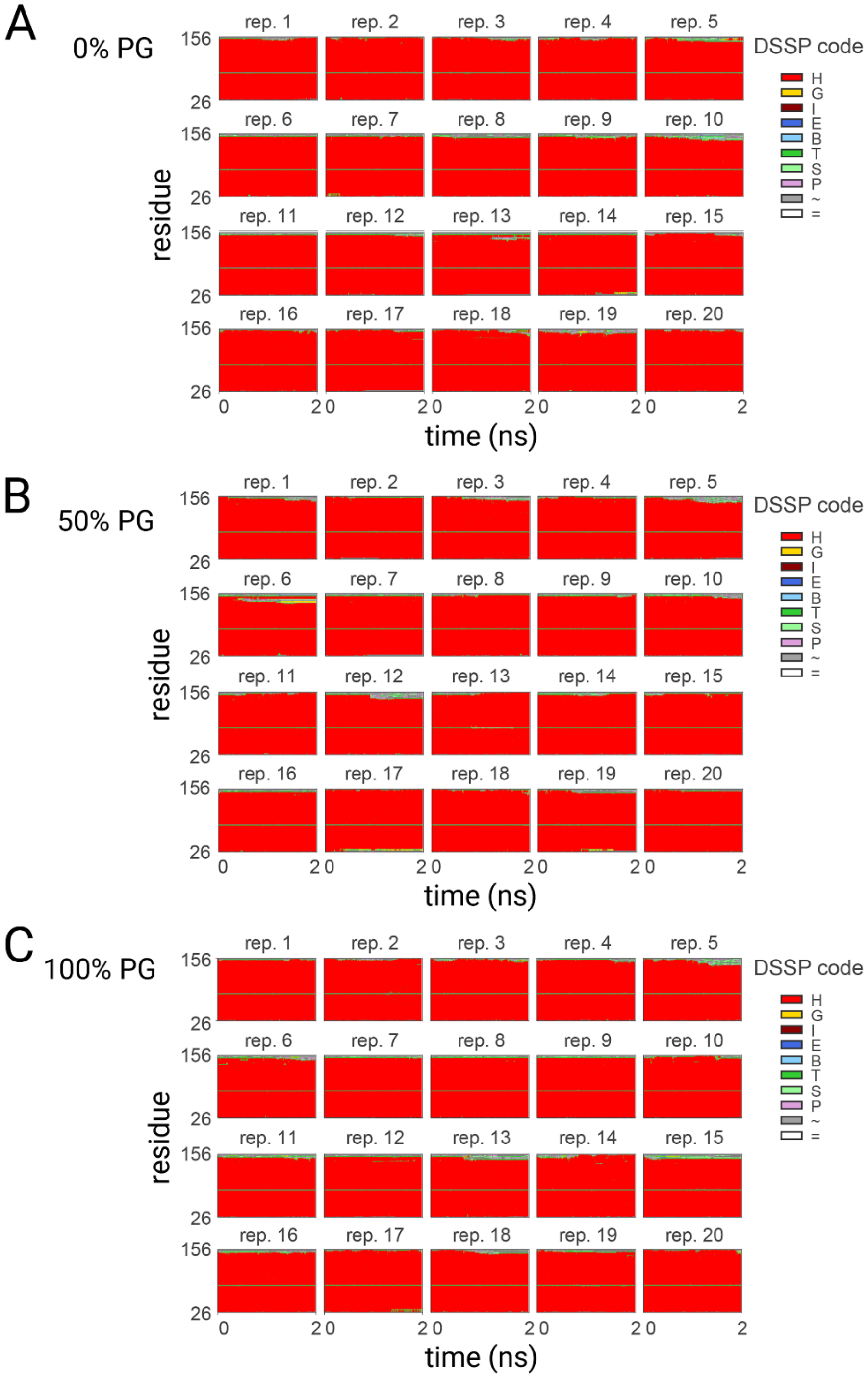


**Figure S8: DSSP analysis of secondary structures of molecular dynamics simulations of IM30 α1-3 in presence of lipid bilayers (heatmap).**

Secondary structure analysis of α1-3 over all replicas across membrane compositions in a time-resolved manner was performed via DSSP to investigate conformational changes during simulations. The x-axes indicate the simulation time of each replicate, while the y-axes display the residues of the peptide. As visible in (A), (B) and (C) the number of c-terminal residues that fluctuate in secondary structure can hereby vary, suggesting that external factors and conformational changes could influence this occurrence, such as binding. In isolated replicas, such as replica 11 of the 50% PG membrane, secondary structure can even be observed in isolated parts of α2/3 around approx. residue 140.

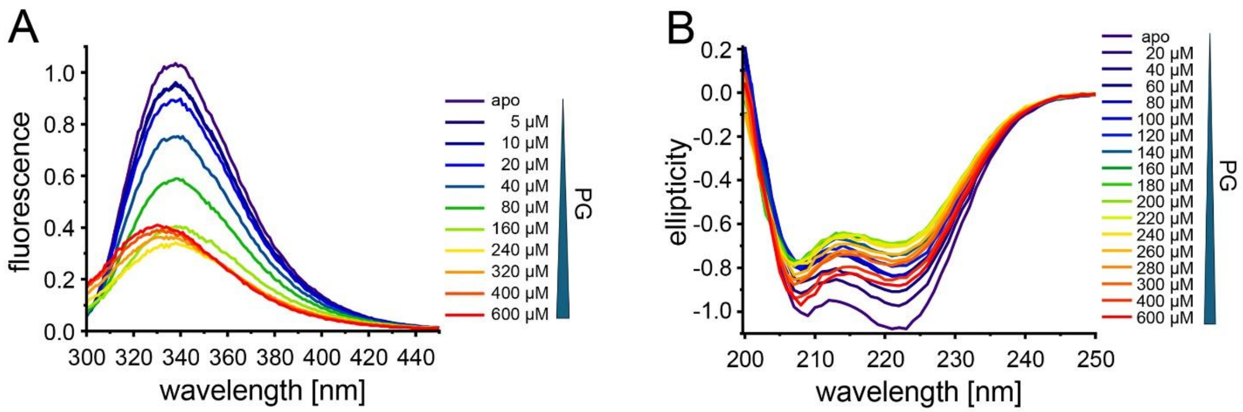


**Figure S9: Membrane binding of IM30 α1-3.**

(A) Representative normalized spectra of the intrinsic protein fluorescence of IM30 α1-3in solution (apo) and after incubation for 2 hours with increasing PG concentrations. The fluorescence emission at 330 nm measured in absence of lipids was set as 1. (B) Representative normalized circular dichroism spectra of IM30 α1-3 in solution (apo) and after incubation for 2 hours with increasing PG concentrations. The ellipticity at 208 nm determined in absence of lipids (apo) was set as -1.0.

**Table S1: Peptides identified via MS.**

Peptide sequences, start and end position of the amino acid residues in the protein sequence of IM30 α1-3, the MaxQuant score, the mass and charge of the peptides, the mass deficit and the number of detected MS/MS spectra are given for each peptide identified in the sample with a score >50.

| Sequence | Start position | End position | Score | Mass | Charges | Mass deficit | MS/MS Count |
|---|---|---|---|---|---|---|---|
| VLEQAVIDMQEDLVQLR | 26 | 42 | 193,69 | 1998,0456 | 2;3 | 0,0865 | 4 |
| LALTNGEENLAR | 75 | 86 | 274,36 | 1299,6783 | 2 | 0,0404 | 4 |
| KSLTDTAAAYQTQLAQQR | 93 | 110 | 146,86 | 1993,0229 | 3 | 0,0661 | 2 |
| SLTDTAAAYQTQLAQQR | 94 | 110 | 161,49 | 1864,9279 | 3 | 0,0300 | 2 |
| TMSENLR | 111 | 117 | 158,53 | 849,40145 | 2 | -0,0293 | 5 |
| RNLAALEAK | 118 | 126 | 154,35 | 984,57163 | 2 | 0,0787 | 3 |
| NLAALEAK | 119 | 126 | 94,806 | 828,47052 | 2 | 0,0494 | 7 |
| ANAELQQTL | 146 | 154 | 91,626 | 986,50327 | 2 | 0,0095 | 2 |
